# Shared inflammatory glial cell signature after brain injury, revealed by spatial, temporal and cell-type-specific profiling of the murine cerebral cortex

**DOI:** 10.1101/2023.02.24.529840

**Authors:** Christina Koupourtidou, Veronika Schwarz, Hananeh Aliee, Simon Frerich, Judith Fischer-Sternjak, Riccardo Bocchi, Tatiana Simon-Ebert, Martin Dichgans, Magdalena Götz, Fabian Theis, Jovica Ninkovic

**Author notes:** these authors equally contributed to the manuscript (Christina Koupourtidou, Veronika Schwarz).

## Abstract

Traumatic brain injury leads to a highly orchestrated immune- and glial cell response partially responsible for long-lasting disability and the development of secondary neurodegenerative diseases. A holistic understanding of the mechanisms controlling the responses of specific cell types and their crosstalk is required to develop an efficient strategy for better regeneration. Here, we combined spatial and single-cell transcriptomics to chart the transcriptomic signature of the injured murine cerebral cortex, and identified specific states of astrocytes, microglia, and oligodendrocyte precursor cells contributing to this signature. Interestingly, these cellular populations share a large fraction of injury-regulated genes, including inflammatory programs downstream of the innate immune-associated pathways Cxcr3 and Tlr1/2. Systemic manipulation of these pathways decreased the reactivity state of glial cells associated with poor regeneration. The functional relevance of the newly discovered shared signature of glial cells highlights the importance of our resource enabling comprehensive analysis of early events after brain injury.

## Introduction

Traumatic brain injury (TBI) defined as acute brain insult due to an external force, such as the direct impact of a penetrating object or acceleration/deceleration force-induced concussions affects people of all ages and is among the major causes of death and disability^1,2^. TBI-induced primary damage leads to neuronal and glial cell death, axonal damage, edema, and disruption of the blood-brain barrier (BBB)^3,4^. The initial insult is followed by progressive secondary damage, which further induces neuronal circuit dysfunction, neuroinflammation, oxidative stress, and protein aggregation. These cellular changes have been associated with prolonged symptom persistence and elevated vulnerability to additional pathologies, including neurodegenerative disorders^4,5^.

TBI-induced pathophysiology evolves through a highly orchestrated response of resident glial cells with peripherally derived infiltrating immune cell populations^3^. After central nervous system (CNS) insult, brain-resident microglia are rapidly activated and change their morphology to a hypertrophic, ameboid morphology^6^. Activated microglia proliferate, polarize, extend their processes, and migrate to the injury site^3,7^. Similarly, oligodendrocyte progenitor cells (OPCs), known as NG2 glia, display rapid cellular changes in response to damage, including hypertrophy, proliferation, polarization, and migration towards the injury site^8–11^. Astrocytes also react to injury with changes in their morphology, gene expression and function in a process referred to as “reactive astrogliosis”^8,12,13^. Reactive astrocytes are characterized by upregulation of intermediate filaments, such as glial fibrillary acidic protein (GFAP), nestin, and vimentin^13–15^. In response to stab-wound injury, astrocytes, in contrast to microglia and OPCs, do not migrate to injury sites, and only a small proportion of astrocytes near blood vessels (juxtavascular astrocytes) proliferate^16^. These initial responses facilitate the formation of a glial border between intact and damaged tissue^12,17,18^, which is necessary not only to restrict the damage^12,17–19^, but also to promote axonal regeneration and circuit restoration^12,19–21^. However, adequate border establishment requires well-orchestrated glial cell reactions in relative distance to the injury site. For example, the distance of astrocytes and OPCs from the injury site has been demonstrated to shape their reactive state^11,16,22^. Furthermore, cross-communication among cell types in several pathological conditions^23–25^, including TBI^26,27^, has been reported to determine cell reactivity states. For example, in neuroinflammatory conditions, reactive microglia induce astrocyte neurotoxicity^28^. Moreover, proliferating astrocytes regulate monocyte invasion^26^, whereas BBB dysfunction alters astrocyte homeostasis and contributes to epileptic episodes^29,30^.

Because the scope of most studies has been restricted to single cellular populations or the interaction of two cell types at most, a detailed investigation of cellular cross-talk after TBI remains lacking. To obtain a holistic understanding of the cellular responses after brain injury, simultaneous examination of multiple cell types in the injury milieu is critical. Therefore, to identify interconnected pathways regulating glial border formation in an unbiased manner, we transcriptomically profiled TBI-induced cell reactivity at spatial and single-cell resolution. Our data provide insights into the spatial, temporal, and single-cell responses of multiple cell types, and reveal a novel, previously overlooked, common injury-induced innate immunity-shared glial signature involving the Toll-like receptor 1/2 (Tlr1/2) and chemokine receptor 3 (Cxcr3) signaling pathways.

## Results

### Brain injury elicits a localized transcriptomic profile in the murine cerebral cortex

TBI induces coordinated cellular reactions leading to glial border formation and isolation of the injury site from adjacent healthy tissue^12^. Importantly, the TBI-induced cellular response is dependent on the distance to the injury site^9,16^. For unbiased identification of regulatory pathways leading to specific spatially defined reactions of glial cells associated with glial border formation, we used spatial transcriptomic (stRNA-seq, i.e., 10x Visium). Stab-wound injuries were induced at the border between the motor and somatosensory cortex in both hemispheres, harming only the gray matter^31^. Because our main focus was on examining the injury-induced changes in the cerebral cortex, we manually resected the mouse brain (Fig. 1a, E.D. Fig. 1a, b). This allowed us to position two parenchymal sections on a single capture area (E.D. Fig. 1a, b). Each section contained the following brain areas: cortex (CTX), white matter (WM), and hippocampal formation (HPF), as identified on the basis of the Allen brain atlas (Fig. 1b).

**Figure 1.**
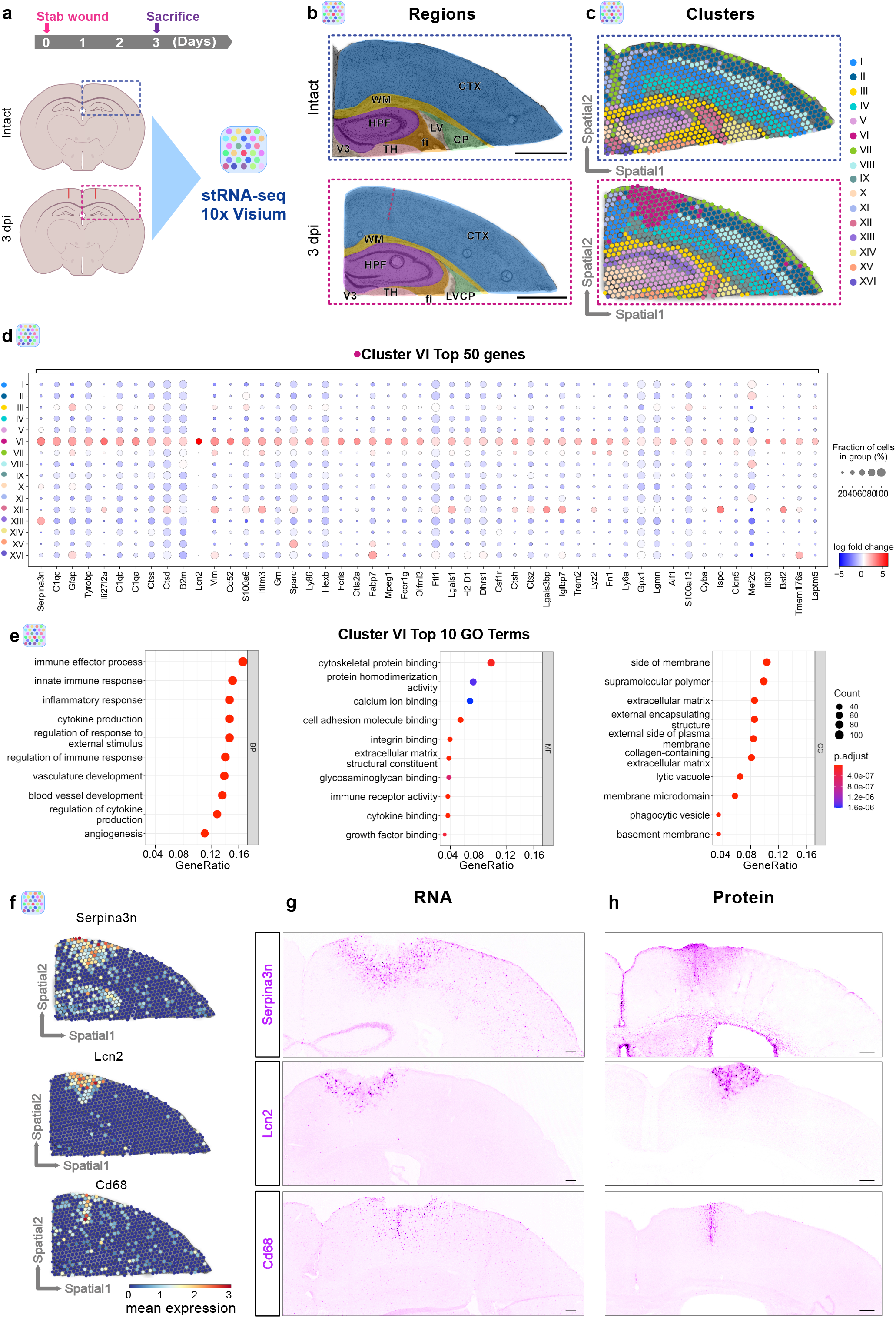
Spatially resolved transcriptomic changes induced by stab wound injury. (**a**) Experimental scheme to conduct spatial transcriptomics in intact and stab wound-injured mouse cerebral cortices (3 dpi). Brains were manually resected, and selected areas highlighted in blue or red dashed boxes were positioned on 10x Visium capture areas. (**b**) Brain sections of both conditions contain cortical, hippocampal, and white matter regions. The dashed red lines indicate the injury cores. (**c**) Clustering of gene expression data on spatial coordinates based on highly variable genes and subsequent dimensionality reduction. (**d**) Dot plot illustrating the expression of the 50 most enriched genes in the injury-induced cluster VI. (**e**) Dot plots depicting GO terms of over-represented cluster VI significantly enriched genes (pval < 0.05, log_2_ fold change > 1). (**f**) Gene expression of cluster VI-enriched genes *Serpina3n, Lcn2*, and *Cd68* in spatial context. (**g**,**h**) Images depicting expression of *Serpina3n, Lcn2*, and *Cd68* at the RNA (**g**) and protein level (**h**) in stab wound-injured cerebral cortices (3 dpi). All images are full z-projections of confocal z-stacks. Scale bars: **g**,**h**: 150 μm. Abbreviations: CTX = cerebral cortex, WM = white matter, HPF = hippocampal formation, LV = lateral ventricle, CP = choroid plexus, V3 = third ventricle, TH = thalamus, fi = fimbria, dpi = days post injury, BP = biological processes, MF = molecular functions, CC = cellular components, GO = gene ontology.

This approach provided the advantage to investigate the expression of a multitude of genes from all cell types at the injury site and to examine their dynamics as a function of distance from the injury site. The primary impact initiates a cascade of processes, which involve the reactions of glial cells and infiltrating or resident immune cells^22,32^. To capture the response of infiltrating immune cells, which peaks at 3 days post-injury (dpi)^26^, and glial cells, which peaks at 2–5 dpi^11,13,26^, we performed stRNA-seq at 3 dpi. Injury-induced alterations were determined by comparison of stab-wounded brain sections to corresponding intact sections (Fig. 1c, E.D. Fig. 1a, b).

Notably, we were able to identify clusters corresponding to specific anatomical structures, e.g., cluster II expressing genes characteristic of cortical layer 2/3 neurons; cluster VIII expressing genes representing layer 4 neurons; and cluster I and cluster IV expressing genes identifying layer 5 and layer 6 neurons, respectively^33^ (E.D. Fig. 1c, d, Ext. Table 2). Importantly, the global cortical layer patterning was not affected by the injury, because we also observed similar gene expression patterns identifying the same neuronal layers in the injured brain sections (E.D. Fig. 1d). However, beyond clusters characterizing individual anatomical structures, we identified an injury-induced cluster, cluster VI, localized around the injury core (Fig. 1b, c, E.D. Fig. 1b). Interestingly, cluster VI was distributed throughout cortical layers 1–5 and was absent in the intact brain sections (Fig. 1c, E.D. Fig. 1a, b). Cluster VI was characterized by a specific transcriptomic signature with enrichment in genes associated with reactive astrocytes^12,34–36^ (*Gfap, Lcn2, Serpina3n, Vim, Lgals1, Fabp7*, and *Tspo*) and microglia^37^ (*Aif1, Csf1r, Cd68*, and *Tspo*), which have been associated with CNS damage^12^ (Fig. 1d, E.D. Fig. 1c, Ext. Table 3).

To obtain insight into the regulated processes within cluster VI, we performed Gene Ontology (GO) enrichment analysis of significantly upregulated genes (pval < 0.05, log_2_ fold change > 1) in this cluster compared with all other clusters. The overrepresented biological processes (BP) were associated with immune response and angiogenesis, whereas the molecular function (MF) and cellular components (CC) indicated changes in genes associated with the extracellular matrix (Fig. 1e, Ext. Table 3). Notably, the above-mentioned processes have been reported to drive glial reaction in response to brain injury and to facilitate glial border formation^19^. Furthermore, processes associated with phagocytosis (lysosome, lytic vacuole, phagocytotic vesicles) were enriched in cluster VI (Fig. 1e, Ext. Table 3), in line with the previously described importance of phagocytotic processes in the context of brain injuries^38^. To confirm the unique injury-induced expression profile and hence the presence of cluster VI, we validated the expression of the cluster VI genes *Serpina3n, Lcn2*, and *Cd68* at the RNA and protein levels. Indeed, the selected candidates were specifically expressed around the injury core, as predicted by our stRNA-seq analysis (Fig. 1f, g), and these expression patterns were also observed at the protein level (Fig. 1h). Although these selected genes were enriched in cluster VI, they displayed unique expression patterns within the injury-induced cluster. Specifically, *Serpina3n* was expressed more broadly than *Lcn2*, whereas *Cd68* displayed a defined expression profile around the injury core (Fig. 1f, g).

To comprehensively determine different expression profiles between the injury core and the perilesional area, we conducted spatial gradient analysis using the SPATA2 analysis pipeline^39^. This allowed us to visualize individual genes and gene set expression patterns as a function of the distance from the injury core (Fig. 2a). For this purpose, we segregated the perilesional area around the injury core into 13 concentric circles (Fig. 2a). In addition, we excluded all subcortical clusters from the analysis and correlated the gene expression profiles along the spatial gradient to a variety of pre-defined models (further details in Methods). To reveal the differences between the injured and perilesional areas, we focused on the genes with “descending” (enriched at the injury core) (Fig. 2b) and “ascending” (depleted at injury core) expression profiles (E.D. Fig. 1e). As expected, all top descending genes were highly enriched at the injury core. However, in the perilesional area (as defined by the border of cluster VI; ∼0.5 mm distance from the injury core) some of these genes displayed unique descending rates (Fig. 2b). We observed heterogeneous expression patterning, ranging from injury core-confined expression (e.g., *Alox5ap* and *Rplp0*) to wide-ranging expression (e.g., *Fth1* and *Gfap*) reaching far from the cluster VI border (Fig. 2c). Of note, gene sets associated with the immune response and inflammation were particularly enriched at the injury core (Fig. 2c), whereas gene sets associated with neuronal and synaptic activity were enriched only in the perilesional areas (E.D. Fig. 1f). Similarly to the descending genes, the ascending genes exhibited relatively divergent expression profiles in the perilesional areas (E.D. Fig. 1e). However, approximately 50% of all top 25 ascending genes were associated with mitochondrial functions, in contrast to the descending genes. These mitochondrial genes exhibited almost identical expression profiles in the perilesional area (E.D. Fig. 1e), thus supporting prior findings that brain insult disrupts normally well-regulated mitochondrial function in a coordinated manner^40–42^.

**Figure 2.**
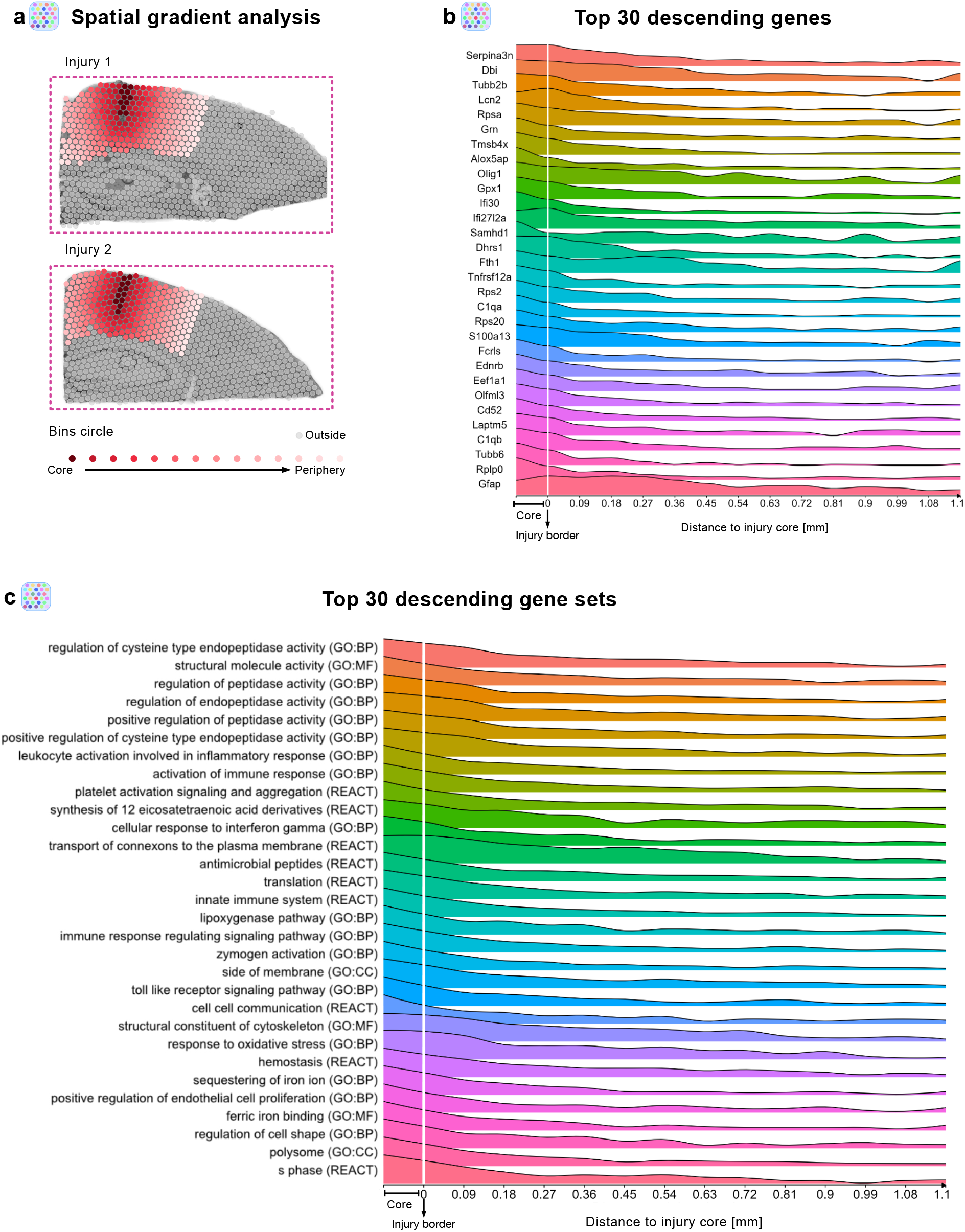
Stab wound injury elicits distinct gene expression patterning along a spatial gradient. (**a**) Paradigm for spatial transcriptomic gradient analysis on stab wound-injured mouse cerebral cortices (3 dpi) by using the SPATA2 pipeline. Spatial gradient analysis was conducted only in cortical areas; from the injury core (dark red spots) toward the periphery (light pink) within 13 concentric circle bins. All other areas (gray spots) were neglected. (**b**) Ridge plot depicting the expression of the 30 most descending genes along the gradient, depicted as mean expression of injury 1 and 2. (**c**) Ridge plot displaying the top 30 most descending gene sets along the gradient, depicted as mean expression from injury 1 and 2. Abbreviations: BP = biological processes, MF = molecular functions, CC = cellular components, GO = gene ontology, REACT = reactome.

In summary, with our spatial gene expression analysis, we identified well-defined anatomical structures as well as an injury-specific cluster characterized by angiogenesis and immune system-associated processes, including phagocytosis. Furthermore, by using spatial gradient analysis, we highlighted injury-induced heterogeneous gene expression profiles in the perilesional area.

### Multiple cellular states contribute to injury-induced local transcriptome profiles

Although stRNA-seq enables profiling of transcriptomic changes by preserving spatial information, the profile itself is derived from multiple cells, which are captured in each spot (1–10 cell resolution). To assess the cellular composition of the injured area and to identify which cell populations defined the transcriptomic profile of cluster VI, we performed single-cell transcriptomic (scRNA-seq) analysis of stab wound-injured cortices and corresponding areas in the intact cortex (3 days post injury, 3 dpi), by using a droplet-based approach (i.e., 10x Chromium) (Fig. 3a). After applying quality control filters, we identified a total of 6322 single cells (Fig. 3b) emerging from both conditions (intact: 2676 cells, 3 dpi: 3646 cells, Fig. 3c, E.D. Fig 2a), which, on the basis of their gene expression, were distributed among 30 distinct clusters. Through this approach, we identified neuronal and glial clusters, including astrocytes, microglia, and oligodendrocyte lineage cells, in addition to vascular cells, pericytes, and multiple types of immune cells (Fig. 3b, E.D. Fig. 2a, b, Ext. Table 5). Additionally, we generated gene expression scores based on established marker genes of well-characterized cell populations in the adult mouse brain (Ext. Table 6). Indeed, the gene scores exhibited enrichment in the corresponding cellular populations, thus further validating our cluster annotation (Fig. 3d).

**Figure 3.**
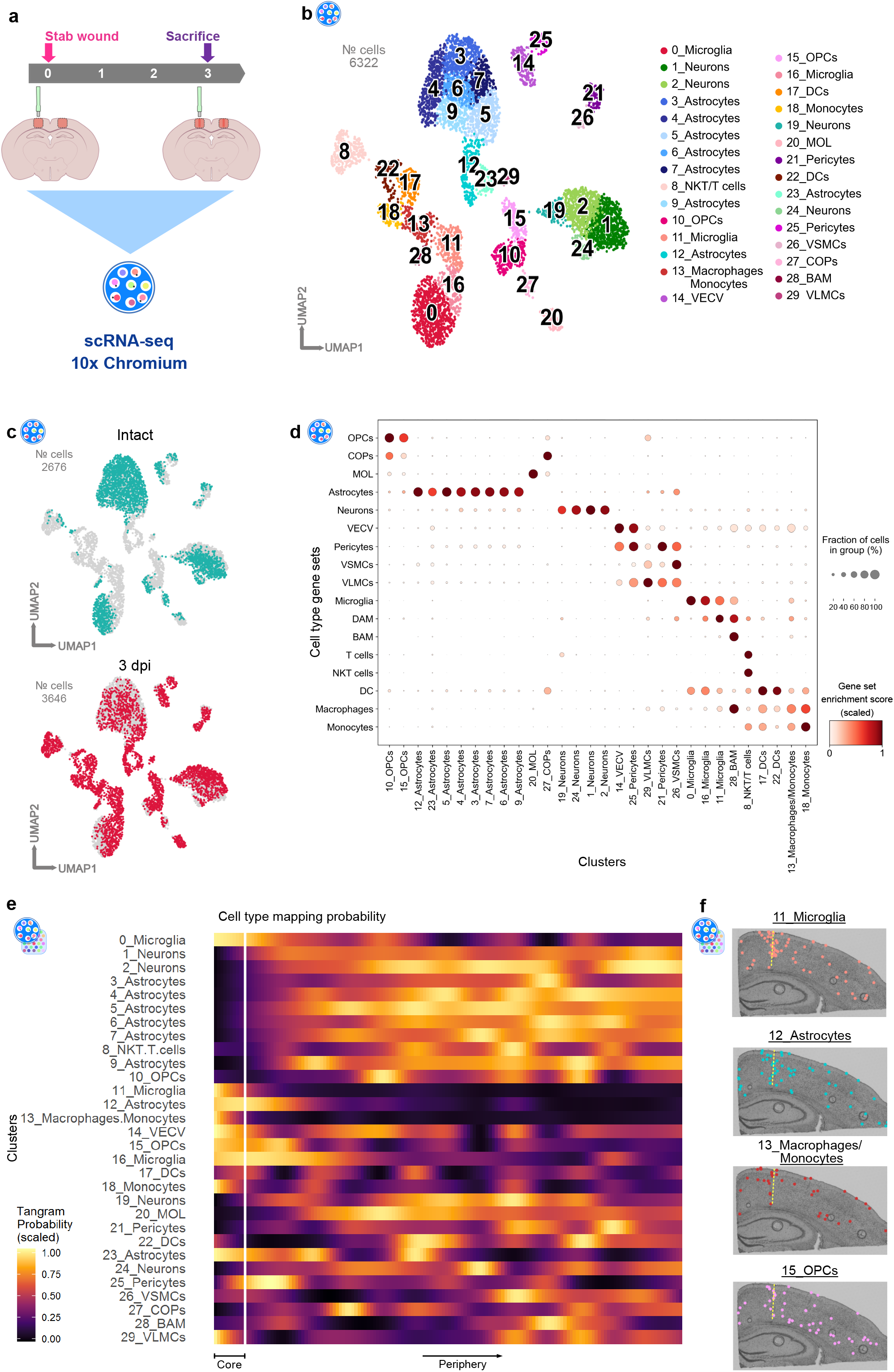
Combination of spatial and single cell transcriptomics identifies cellular populations contributing to distinct transcriptional responses at the injury site. (**a**) Experimental scheme to conduct single-cell RNA-sequencing of intact and stab wound-injured cerebral cortices (3 dpi) with the 10x Genomics platform. Red masked areas on brain schemes indicate biopsy areas used for the analysis. (**b**) UMAP plot illustrating 6322 single cells distributed among 30 distinct clusters. Clusters are color-coded and annotated according to their transcriptional identities. (**c**) UMAP plot depicting the distribution of cells isolated from intact (green) and injured (red) cerebral cortices. (**d**) Dot plot indicating strong correlation of post hoc cluster annotation with established cell type-specific gene sets (Ext. Table 6). (**e, f**) 3 dpi scRNA-seq cluster localization along the spatial gradient (Fig. 2), based on probabilistic mapping with Tangram (**e**) and single cell deconvolution (**f**) in a spatial context. Abbreviations: UMAP = uniform manifold approximation and projection, dpi = days post injury, OPCs = oligodendrocyte progenitor cells, COPs = committed oligodendrocyte progenitors, MOL = mature oligodendrocytes, VECV = vascular endothelial cells (venous), VSMCs = vascular smooth muscle cells, VLMCs = vascular and leptomeningeal cells, DAM = disease-associated microglia, BAM = border-associated macrophages, NKT cells = natural killer T cells, DC/DCs = dendritic cells.

Interestingly, by comparing the cell distributions between the intact and injured conditions, we observed that several clusters of immune and glial cells were highly abundant exclusively in the injured brain (Fig. 3c, E.D. Fig. 2a). The clusters 8_NKT/T cells, 13_Macrophages/Monocytes, 17_DCs, 18_Monocytes, and 22_DCs, for example, appeared primarily after injury and expressed *Ccr2*^6,22^ (E.D. Fig. 2c) in addition to their distinct cell identity markers (E.D. Fig. 2b, Ext. Table 5). Microglia clusters that appeared after injury (11_Microglia and 16_Microglia) exhibited high expression of *Aif1* and low expression of the homeostatic microglia markers *Tmem119* and *P2ry12*^43^ (E.D. Fig. 2c). Similarly, the astrocytic clusters 12_Astrocytes and 23_Astrocytes were present primarily in the injured condition and were characterized by high expression of classical reactive astrocyte markers such as *Gfap* and *Lcn2*^34,36^ (E.D. Fig. 2d). In addition to microglia and astrocytes, the cluster 15_OPCs was present primarily after injury (E.D. Fig. 2a). Cells from cluster 15_OPCs expressed a combination of genes associated with the cell cycle (G2/M phase, E.D. Fig 2f, Ext. Table 7^44,45^) and *Cspg4* (E.D. Fig. 2e); both hallmarks of Nerve/glial antigen 2 glia (NG2 glia), which rapidly proliferate after brain injury^9^.

To elucidate which of these cellular clusters contributed to the injury-specific signature of cluster VI, we mapped the single cell expression data onto the spatial gene expression dataset (Fig. 3e, f, E.D. Fig. 3 and 4) by using Tangram^46^. To include the identical anatomical regions regarding the scRNA-seq data acquisition, we restricted the stRNA-seq dataset to the cortical clusters (clusters I, II, IV, VI, VII, VIII, and IX). The probabilistic mapping predicted that several clusters including 11_Microglia, 16_Microglia, 12_Astrocytes, 23_Astrocytes, 13_Macrophages/Monocytes, 18_Monocytes, and 15_OPCs were localized near the injury core (Fig. 3e, E.D. Fig. 3a). In contrast, neuronal clusters 1_Neurons, 2_Neurons, and 24_Neurons, as well as the astrocytic clusters 3_Astrocytes, 5_Astrocytes, 7_Astrocytes, and 9_Astrocytes, displayed decreased representation around the injury site (Fig. 3e, E.D. Fig. 3a). Additionally, we used the H&E images of the stRNA-seq dataset to estimate the number of nuclei within each spot of the capture area, which, in combination with probabilistic mapping, can be used for deconvolution. To some extent, this analysis further associated the above-mentioned clusters with the injury milieu (Fig. 3f, E.D. Fig. 4a, b). Importantly, not all glial cells contributed to the injury environment (E.D. Fig. 3a, 4b). Most astrocytic clusters, with the exception of clusters 12_Astrocytes and 23_Astrocytes, did not show enriched mapping at the injury site (Fig. 3e, E.D. Fig. 3a). Similar behavior was detected for the oligodendrocyte clusters 20_MOL and 27_COPs (E.D. Fig. 3a). Notably, our deconvolution analysis supported these observations (E.D. Fig. 4b). In summary, the combination of stRNA-seq with corresponding scRNA-seq datasets allowed us to identify an injury-specific transcriptional profile exhibiting enrichment of individual glial subpopulations, and subsequent depletion of distinct astrocytic and neuronal clusters.

### Injury induces common transcriptomic changes in glial cells

Because glial cell reactivity exhibits distinct temporal dynamics in response to injury^11,13,26^, we decided to add an additional time point (5 dpi) to our scRNA-seq analysis (Fig. 4a). This experimental design enabled investigation of the transcriptional states of glial cells underlying the observed heterogeneity in glial cell responses. In total, we analyzed 33862 cells (intact: 16567, 3 dpi: 3637, 5 dpi: 13658), which were distributed among 35 clusters (Fig. 4b, E.D. Fig. 5a-c, Ext. Table 8). In line with our previous observations (Fig. 3c), a comparison of cell distribution among all three conditions (intact, 3 dpi, and 5 dpi) elucidated several injury-induced clusters, which were formed exclusively by cells originating from brain-injured animals (E.D. Fig. 5b). However, none of these injury-induced clusters were specific to either the 3 or 5 dpi time point (E.D. Fig. 5b).

**Figure 4.**
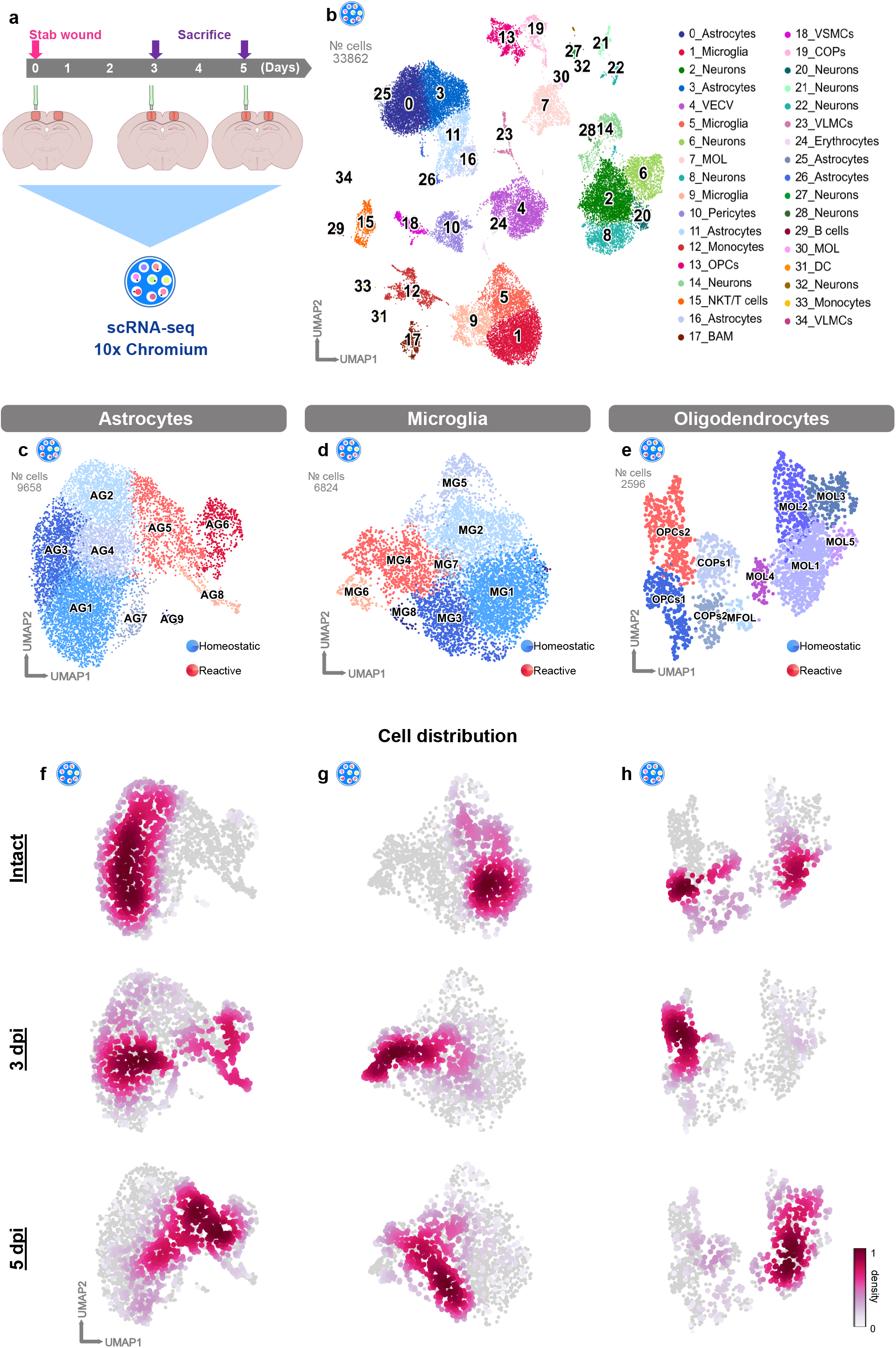
Stab wound injury induces defined transcriptional changes in glial subpopulations. (**a**) Experimental scheme for single-cell RNA-sequencing of intact and stab wound-injured cerebral cortices (3 and 5 dpi) with the 10x Chromium platform. Red masked areas on brain schemes indicate biopsy areas used for the analysis. (**b**) UMAP embedding of integrated and batch-corrected single cell transcriptomes of 33862 cells distributed among 35 distinct clusters. Clusters were color-coded and annotated on the basis of their transcriptional identities. (**c-e**) UMAPs depicting subclustering of astrocytes (9 clusters) (**c**), microglia (8 clusters) (**d**), and oligodendrocytes (10 clusters) (**e**). Cells were further assigned to homeostatic (blue) or reactive (red) clusters according to cell origin (E.D. Fig. 6) and distinct marker expression. (**f-h**) UMAPs illustrating cell distributions at all time points (intact, 3 dpi, and 5 dpi) among subclusters of astrocytes (**f**), microglia (**g**), and oligodendrocytes (**h**). Data were downsampled to an equal number of cells between time points for each cell type (**f-h**). Abbreviations: UMAP = uniform manifold approximation and projection, dpi = days post injury, OPCs = oligodendrocyte progenitor cells, COPs = committed oligodendrocyte progenitors, MOL = mature oligodendrocytes, VECV = vascular endothelial cells (venous), VSMCs = vascular smooth muscle cells, VLMCs = vascular and leptomeningeal cells, BAM = border-associated macrophages, NKT cells = natural killer T cells, DC/DCs = dendritic cells.

To unravel how each population transited from a homeostatic to a reactive state, we focused on individual cell populations. Hence, we further subclustered astrocytes, microglia, and oligodendrocytes (Fig. 4c-e, E.D. Fig. 6). We identified distinct clusters in each of the investigated populations, which were composed primarily of cells from intact (blue clusters) and injured (orange/red clusters) samples (Fig. 4c-e, E.D. Fig. 6c, h, m). Additionally, cells originating from the injured samples expressed typical markers of glial reactivity. We identified clusters AG5, AG6, and AG8 as the main populations of reactive astrocytes (Fig. 4c, E.D. Fig. 6a-e), because these cells expressed high levels of *Gfap, Vim*, and *Lcn2*^34,36^ (E.D. Fig. 6e). Microglial clusters MG4 and MG6 displayed high expression of *Aif1* and low expression of the homeostatic markers *Tmem119* and *P2ry12*^43^(Fig. 4d, E.D. Fig. 6f-j). By subclustering cells belonging to the oligodendrocytic lineage, we were able to identify two populations of OPCs (OPCs1 and OPCs2) (Fig. 4e, E.D. Fig. 6k-o). Cluster OPCs2 was composed primarily of cells from injured samples (Fig. 4e). Of note, we were not able to find a unique marker within the OPCs2 cluster for identifying reactive OPCs (E.D. Fig. 6o). Importantly, cells from the injury-responding clusters, as identified by our previous deconvolution analysis (11_Microglia, 12_Astrocytes, and 15_OPCs) (Fig. 3f, E.D. Fig. 4), also mapped predominantly to the glial subclusters evoked by injury (E.D. Fig. 6d, i, n). Together, these results corroborated the reactive state of these glial subclusters (henceforth referred to as reactive clusters).

By subclustering glial cells, we did not discover any cluster unique to either the 3 or 5 dpi time point. This finding suggests a gradual activation of glial cells in response to injury rather than distinct activation states. To shed light on this possibility, we examined the cell distributions of all subclustered glial cells among all time points (intact, 3 dpi, and 5 dpi) (Fig. 4f-h). Interestingly, we observed prominent changes in the distribution of reactive clusters between time points after injury. More specifically, many of the astrocytes at 3 dpi remained present in the homeostatic clusters and were only partially present in the reactive clusters AG6 and AG8, whereas at 5 dpi, most cells were detected in cluster AG5 (Fig. 4f). Microglia, in contrast, displayed a faster transition to reactivity than astrocytes: at 3 dpi, most cells were already localized in the reactive clusters MG4 and MG6. At 5 dpi, however, most of the cells had begun to transition back to the homeostatic state, and a high proportion of cells were present in cluster MG3 (Fig. 4g). Similarly, OPCs reacted rapidly after injury, because at 3 dpi, most cells resided in the reactive cluster OPCs2, whereas only several cells were present in this cluster at 5 dpi (Fig. 4h). These results are in line with prior findings showing that microglia and OPCs rapidly respond to injury, and their reactivity peak ranges from 2 to 3 dpi, whereas astrocyte reactivity peaks at 5 dpi^11,13,26^.

The enriched immune system-associated processes around the injury site, as indicated by spatial transcriptomics, and the identification of specific reactive subtypes of glial cells populating the injury environment, prompted the question of whether the inflammatory gene expression might be a unique signature of one specific cell type or a common feature of reactive glia. Therefore, within each glial population (Fig. 4 c-e), we extracted the differentially expressed genes (DEGs) of each glial subcluster (pval < 0.05, log_2_ fold change > 1.6 or < -1.6) and compared them among all subclusters (E.D. Fig. 7a, b, Ext. Table 9). Interestingly, among all clusters, the highest similarity of upregulated genes was observed between the reactive glial clusters MG4, AG5, and OPCs2, with 66 enriched genes in common (E.D. Fig. 7b). This finding suggested that, in response to injury, individual reactive glial clusters might share some cellular programs. Hence, we performed GO term analysis of the 192 commonly upregulated genes from the comparison of clusters MG4, AG5, and OPCs2, independently of other glial subclusters (Fig. 5a). Most of the commonly regulated processes were associated with cell proliferation (Fig. 5b, Ext. Table 10), which has been reported to be a shared hallmark of glial cell reactivity^13,34^. Moreover, we identified processes associated with innate immunity (Fig. 5b, Ext. Table 10) and numerous genes associated with the type I interferon signaling pathway (*Ifitm3, Ifit3, Bst2, Isg15, Ifit3b, Irf7, Ifit1, Ifi27l2a, Oasl2*, and *Oas1a*) (Fig. 5c) as well as *Cxcl10*, a ligand activating the Cxcr3 pathway^47^. Indeed, by using RNA scope, we confirmed that *Cxcl10, Oasl2*, and *Ifi27l2a* were expressed by a subset of microglia, astrocytes, and OPCs (Fig. 5d-g, E.D. Fig. 8a-g). Furthermore, we observed shared expression of Galectin1 (*Lgals1*) by a subset of microglia, astrocytes, and OPCs at the protein level (E.D. Fig. 9a-d). Notably, the expression of these innate immunity-associated genes was clearly restricted to distinct glial subpopulations, because not all glial cells expressed these markers (Fig. 5d-g, E.D. Fig. 8a-g, E.D. Fig. 9a-d). Additionally, the upregulation of the innate immunity-associated genes (Fig. 5c) after injury is a unique feature of glial cells because neurons never expressed these genes, whereas in vascular cells they were present at low levels in the intact brain (E.D. Fig. 9e, f).

**Figure 5.**
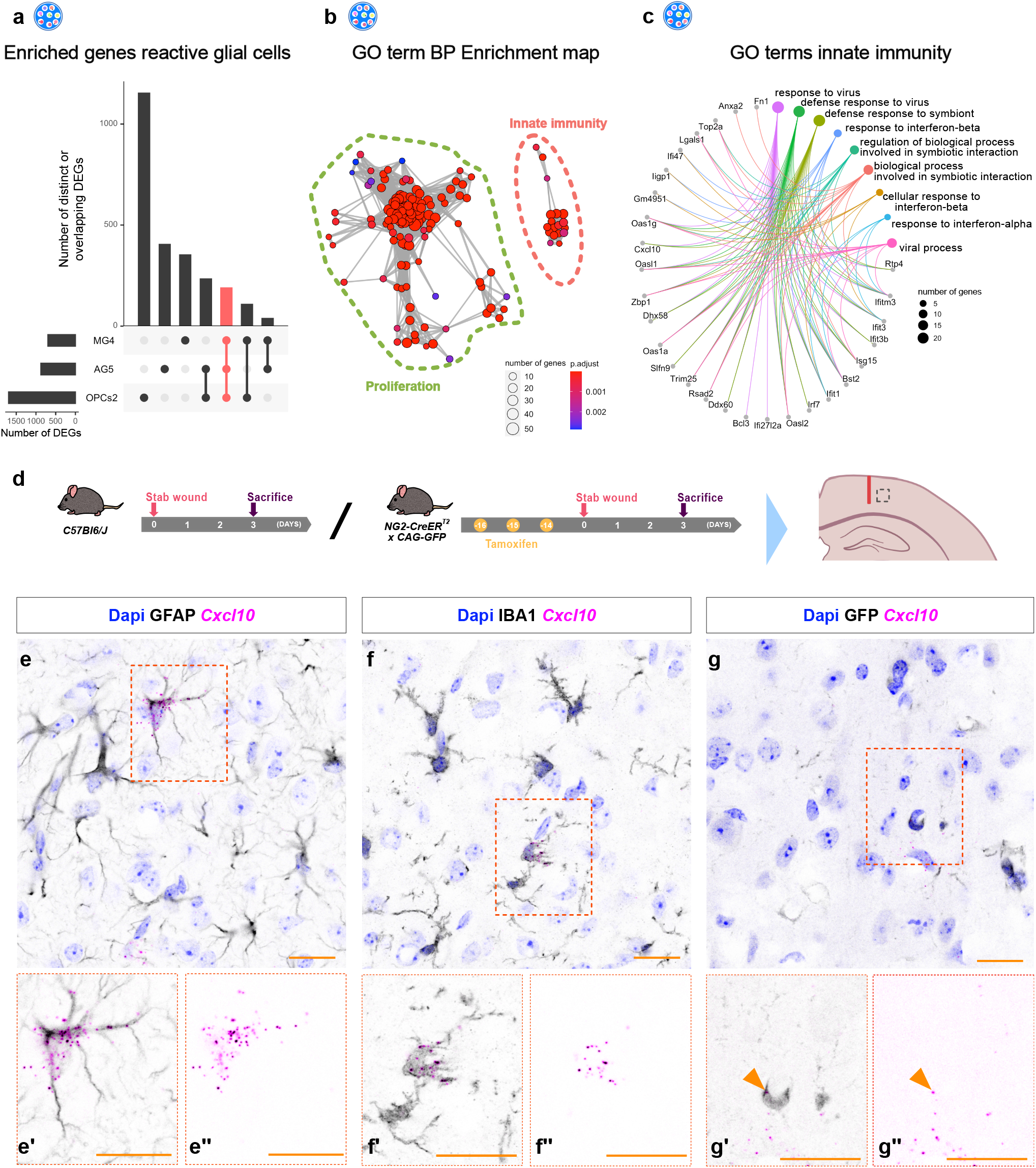
Stab wound injury induces common transcriptional changes in reactive glial subpopulations. (**a**) UpSet plot depicting unique (single points) or overlapping (connected points) DEGs (pval < 0.05, log_2_ fold change > 1.6) induced in reactive glial subclusters MG4, AG5, and OPCs2, compared with all other clusters of the respective cell type. A total of 192 commonly shared genes between these clusters are highlighted by the red bar. (**b**) GO term network analysis of the 192 commonly shared genes, associating shared DEGs with the biological processes of proliferation (green dashed line) and innate immunity (orange dashed line). (**c**) Chord diagram illustrating innate immunity GO terms from Fig. 5b and the corresponding genes. (**d**) Experimental paradigm for visualizing shared gene expression in astrocytes and microglia (*C57Bl6/J*) as well as OPCs (*NG2-CreER*^*T2*^*xCAG-GFP*). The dashed gray box on the mouse brain scheme refers to an example imaging area. The red line displays injury core. (**e-g**) RNAscope example images of the shared innate immunity-associated gene *Cxcl10* (magenta) counterstained with GFAP (black) (**e**), IBA1 (black) (**f**), and GFP (NG2^+^ glia) (black) (**g**) antibodies. Micrographs (**e’-g’**) are magnifications of the red boxed areas in the corresponding overview images. Orange arrowheads in micrograph (**g’-g”**) indicate colocalization of *Cxcl10* with GFP^+^ cells. All images are single z-plane of confocal z-stacks. Scale bars: (**e-g’’**): 20 μm. Abbreviations: DEGs = differentially expressed genes, GO = gene ontology, BP = biological processes.

We further asked whether the shared inflammatory signature might be a conserved feature of reactive glia in different pathological conditions. We used a publicly available database describing the response of astrocytes to systemic lipopolysaccharide (LPS) injection.^36^ Interestingly, a high proportion of the shared inflammatory marker genes (e.g., *Oasl2* and *Ifit1*) were expressed exclusively in the astrocytic cluster 8 (E.D. Fig. 10a). Astrocytes of cluster 8 were classified in the above-mentioned study as reactive astrocytes in a sub-state capable of rapidly responding to inflammation. Notably, not all inflammatory genes were detected in LPS-induced reactive astrocytes, thus indicating that only a portion of the signature was retained (E.D. Fig. 10b). Furthermore, we investigated whether human iPSC-derived reactive astrocytes displayed similar inflammatory responses to those in stab wound-injured mice. Therefore, we mapped the genes characterizing the two inflammatory clusters of iPSC-derived reactive astrocytes from Leng et al.^48^ (IRAS 1 and IRAS2) on our integrated single-cell dataset (Fig. 4b, E.D. Fig. 10c).^49^ Interestingly, the inflammatory signatures of both iPSC-derived reactive astrocyte clusters (IRAS 1 and IRAS2) were induced primarily in the subclusters of astrocytes that emerged after injury (11_Astrocytes and 16_Astrocytes) (E.D. Fig. 5a, b, E.D. Fig. 10c, d), thus confirming common reactive astrocytic states between murine and human glia. Of note, the inflammatory signature of the human iPSC-derived astrocytes was not observed exclusively in reactive astrocytes but was highly abundant in other cellular populations (E.D. Fig. 10c, d). This finding strongly emphasizes the need for a holistic cellular view of brain pathologies to identify therapeutical targetable pathways. Our findings revealed a common inflammatory signature present in a subset of reactive glial cells in response to TBI. Moreover, this shared inflammatory signature is largely preserved in different pathological conditions and species.

### Regulation of injury-induced innate immune responses via the Cxcr3 and Tlr1/2 pathways

On the basis of the shared regulation of the innate immunity pathways after brain injury, including components of the CXC chemokine receptor 3 (Cxcr3) and Toll-like receptor 2 (Tlr2) pathways (E.D. Fig. 9g), and our recent findings that Cxcr3 and Tlr1/2 regulate OPC accumulation at injury site in the zebrafish brain^49^, we investigated the injury-induced transcriptional changes after interference with the Cxcr3 and Tlr1/2 pathways. We used the Cxcr3 antagonist NBI-74330^50^ and the Tlr1/2 pathway inhibitor CU CPT 22^51^ to interfere with the above-mentioned pathways, and performed scRNA-seq analysis at 3 dpi and 5 dpi (henceforth referred to as SW INH or 3/5 dpi_INH if distinct time points are indicated) (Fig. 6a). The specificity of these chemical compounds was validated with a murine knock-out OPC cell line^49^. The data were integrated with our previously acquired datasets (Fig. 4b) from intact (INT) and injured animals (henceforth referred to SW CTRL or as 3/5 dpi_CTRL if distinct time points are indicated). In total, we analyzed 53813 cells (INT: 16649 cells, 3 dpi_CTRL: 3643 cells, 3 dpi_INH: 4613 cells, 5 dpi_CTRL: 13766 cells, 5 dpi_INH: 15142) were distributed among 36 clusters (Fig. 6b, E.D. Fig. 11a). Notably, with the integration of additional conditions (3 and 5 dpi_INH), and hence a subsequent increase in the total cell number, we did not observe the emergence of new clusters (E.D. Fig. 11a). Furthermore, even after the integration of SW INH datasets, the overall cluster identity was unaffected, as indicated by high similarity scores among the clusters (E.D. Fig. 11b). Because microglia, astrocytes, and OPCs displayed common innate immune-associated gene expression after stab wound injury (Fig. 5c-g), we sought to investigate the possible influence of Cxcr3 and Tlr1/2 pathway inhibition on microglia, astrocytes, and OPCs by further subclustering the above-mentioned cell types. In each investigated cell population, we again identified distinct clusters containing primarily cells from injured samples (Fig. 6c-e, E.D. Fig. 11c-e). Of note, these clusters were composed of cells originating from both SW CTRL and SW INH samples. These results suggested that the inhibition of Cxcr3 and Tlr1/2 pathways after stab-wound injury did not induce new transcriptional states. Instead, the inhibitor treatment resulted in partial downregulation of the inflammatory genes (Fig. 5c) shared among the reactive clusters AG7, MG3, and OPCs2 (Fig. 6f, Ext. Table 11).

**Figure 6.**
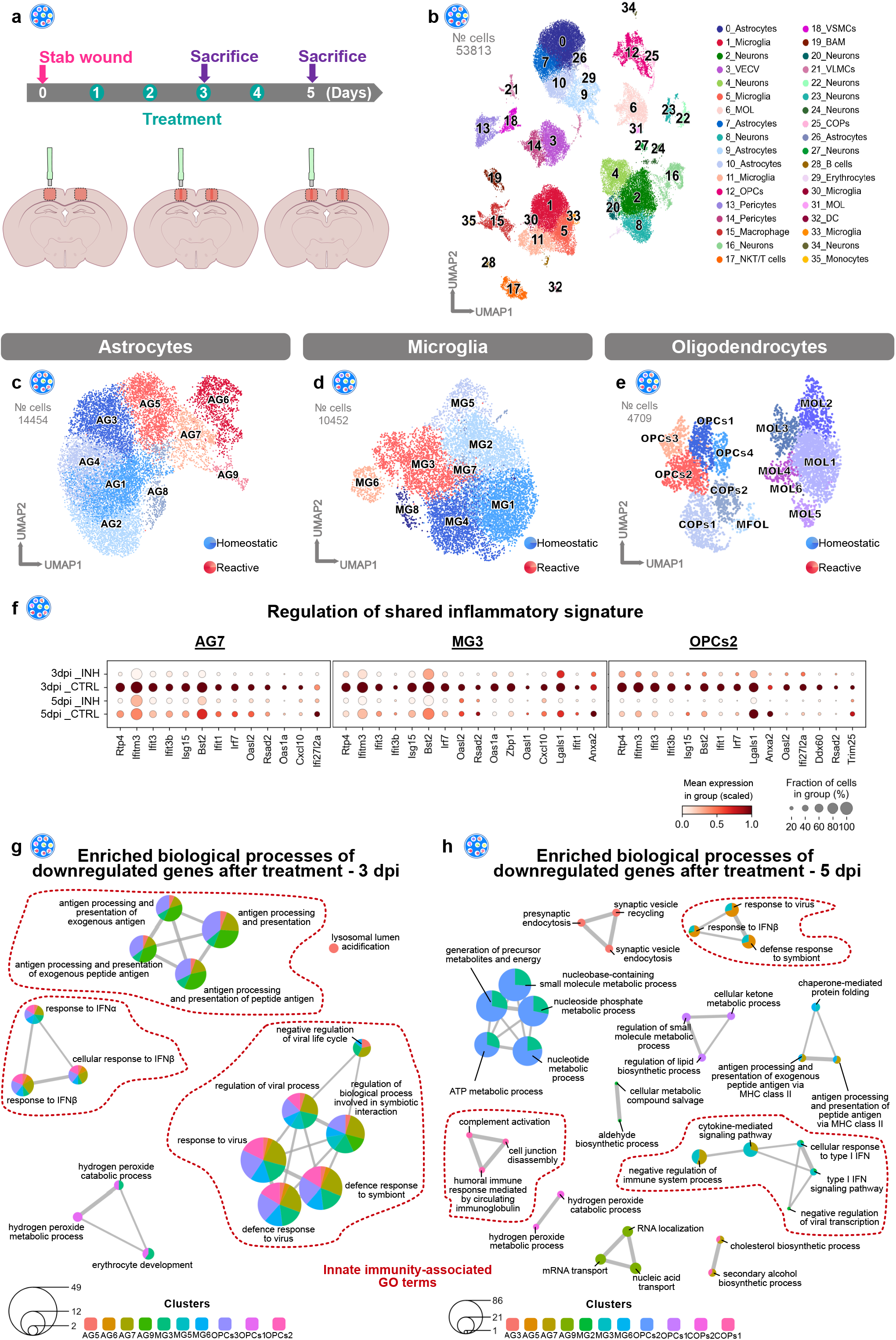
The Cxcr3 and Tlr1/2 pathways orchestrate the innate immune response shared among reactive glial cells. (**a**) Experimental scheme for single-cell RNA-sequencing of intact, stab wound-injured control (3/5 dpi_CTRL) and stab wound-injured inhibitor-treated (3/5 dpi_INH) cerebral cortices with the 10x Chromium platform. Red masked areas on brain schemes indicate biopsy areas used for the analysis. (**b**) UMAP embedding of integrated and batch-corrected single cell transcriptomes of 53813 cells. Cells were distributed among 36 distinct clusters, color-coded, and annotated according to their transcriptional identities. (**c-e**) UMAPs illustrating subclustering of astrocytes (9 clusters) (**c**), microglia (8 clusters) (**d**) and oligodendrocytes (13 clusters) (**e**). Cells were further assigned to homeostatic (blue), or reactive (red) clusters according to cell origin (E.D. Fig. 10). (**f**) Dot plots depict decreased expression of various shared inflammatory genes (Fig. 4c) in the reactive glial clusters AG7, MG3, and OPCs2 after inhibitor treatment. (**g-h**) GO term networks illustrating common and unique downregulated biological processes of glial subclusters in response to Cxcr3 and Tlr1/2 pathway inhibition at 3 dpi (**g**) and 5 dpi (**h**) (Ext. Table 12). Common downregulation of innate immunity-associated GO terms (highlighted by red dashed lines) are observed at 3 but not at 5 dpi. Abbreviations: UMAP = uniform manifold approximation and projection, dpi = days post injury, OPCs = oligodendrocyte progenitor cells, COPs = committed oligodendrocyte progenitors, MOL = mature oligodendrocytes, VECV = vascular endothelial cells (venous), VSMCs = vascular smooth muscle cells, VLMCs = vascular and leptomeningeal cells, BAM = border-associated macrophages, NKT cells = natural killer T cells, DC/DCs = dendritic cells.

To address transcriptional changes induced by the inhibitor treatment, we performed differential gene expression analysis of each subcluster between SW CTRL and SW INH at each time point (pval < 0.05, log_2_ fold change > 0.7 or log_2_ fold change < -0.7). Interestingly, most of the inhibitor-induced changes at 3 and 5 dpi were subcluster specific, because only a few DEGs overlapped (E.D. Fig. 12a-d). To reveal the biological processes regulated in each glial subcluster (Fig. 6c-e), we used the function compareCluster^52^ (clusterProfiler R package) and calculated the enriched functional profiles of each cluster. This function summarized the results into a single object and allowed us to compare the enriched biological processes of all glial subclusters at once. Indeed, by comparing the processes associated with all significantly downregulated genes after treatment at 3 dpi, we identified many programs associated with the innate immune response, which were shared among several glial populations, including reactive astrocytes (clusters AG5, AG6, AG7, and AG9), microglia (clusters MG3 and MG6), and OPCs (cluster OPCs2) (Fig. 6g, Ext. Table 12). Interestingly, although still downregulated at 5 dpi, these immune response-associated processes were no longer shared among different glial populations (Fig. 6h, Ext. Table 12). In contrast, biological processes induced by the inhibitor treatment were cluster specific, independently of the analysis time point (E.D. Fig. 12e, f). Together, our scRNA-seq analysis findings indicated that the Cxcr3 and Tlr1/2 signaling pathways regulate similar processes in initial activation (3 dpi) of different glial cells. However, this activation is followed by cell type-specific transcriptional changes at later stages (5 dpi).

### Inhibition of the Cxcr3 and Tlr1/2 signaling pathways does not interfere with oligodendrocyte reactivity and proliferation

Interference with the Cxcr3 and Tlr1/2 signaling pathways after brain injury did not result in the emergence of new cell types or states at either 3 or 5 dpi (E.D. Fig. 11a). Nevertheless, the inhibition of the above-mentioned pathways elicited an overall downregulation of various inflammation-associated genes in the reactive glial clusters AG7, MG3, and OPCs2, particularly at 3 dpi (Fig. 6f, Ext. Table 11). Furthermore, inhibition of the Cxcr3 and Tlr1/2 pathways after injury in the zebrafish telencephalon modulated oligodendrocyte proliferation, thereby decreasing oligodendrocytes in the injury vicinity^49^. To investigate the relevance of Cxcr3 and Tlr1/2 signaling in the mammalian context, we first sought to examine the cluster distribution of oligodendroglial lineage cells among all conditions (INT, SW CTRL and SW INH) and time points (3 and 5 dpi) (E.D. Fig. 13a). Surprisingly, we detected no differences in the cluster distribution of the reactive OPC clusters OPCs2 and OPCs3 between SW CTRL and SW INH cells at 3 or 5 dpi (E.D. Fig. 13b). To further corroborate our scRNA-seq findings, we determined the number of OLIG2^+^ oligodendrocytes in stab wound-injured mice at 3 dpi (E.D. Fig. 13c). In line with the findings from our computational analysis, we detected no differences in the number of OLIG2^+^ cells near the injury site between the experimental groups (E.D. Fig. 13d-f). Finally, we determined the proliferation ability of OLIG2^+^ cells between both experimental groups by labeling all cells in S-phase with the DNA base analogue EdU (0.05 mg/g 5-ethinyl-2′-deoxyuridine i.p. injection 1 hr before sacrifice) and observed no changes in the number of proliferating (OLIG2^+^ and EdU^+^) oligodendrocytes (E.D. Fig. 13d, e, g).

In summary, the inhibition of Cxcr3 and Tlr1/2 signaling pathways after stab wound injury in the mouse cerebral cortex, in contrast to findings in the zebrafish brain, did not alter oligodendrocyte proliferation or affected the overall number of oligodendrocyte lineage cells near the injury site early after injury, but did alter their inflammatory signatures.

### The Cxcr3 and Tlr1/2 signaling pathways regulate microglial activation in response to injury

The expression of inflammatory genes in microglia is tightly associated with their activation^7,53^. Therefore, we assessed whether the downregulation of inflammatory genes induced by Cxcr3 and Tlr1/2 inhibition (Fig. 6f) might alleviate microglial reactivity. Hence, we examined the cell distribution of subclustered microglia among all three conditions (INT, SW CTRL, and SW INH) and time points (3 and 5 dpi) (Fig. 7a, b). As previously depicted in Fig. 4g, cells derived from intact samples were confined to the homeostatic clusters, whereas cells from the injured samples were distributed primarily in the reactive clusters at 3 dpi, and a transition toward the homeostatic clusters was noticeable at 5 dpi. A direct comparison of SW CTRL and SW INH samples indicated differences in the cell distributions, with a higher proportion of cells localized in the homeostatic clusters after Cxcr3 and Tlr1/2 inhibition (Fig. 7b). Although the discrepancy between conditions was already detectable at 3 dpi, the shift was more pronounced at 5 dpi (Fig. 7b). To further elucidate whether the detected shift in microglia distribution after Cxcr3 and Tlr1/2 pathway inhibition was accompanied by changes in overall cell morphology, we assessed microglia cell characteristics with the automated morphological analysis tool described by Heindl et al.^54^. Brain sections from SW CTRL and SW INH animals were labeled with an anti-IBA1 antibody, and areas near the injury site were analyzed (Fig. 7c). Microglia from SW INH brains displayed significantly smaller cell somata, a less round shape, and greater branch length than microglia from SW CTRL brains (Fig. 7d-f, E.D. Fig. 14a). The inhibition of Cxcr3 and Tlr1/2 signaling pathways decreased branch volume without altering the total number of major branches (E.D. Fig. 14a). In addition, although not significantly altered, microglia from SW INH brains appeared to be more ramified than SW CTRL microglia, because more nodes per major branch were detected. (E.D. Fig 14a).

**Figure 7.**
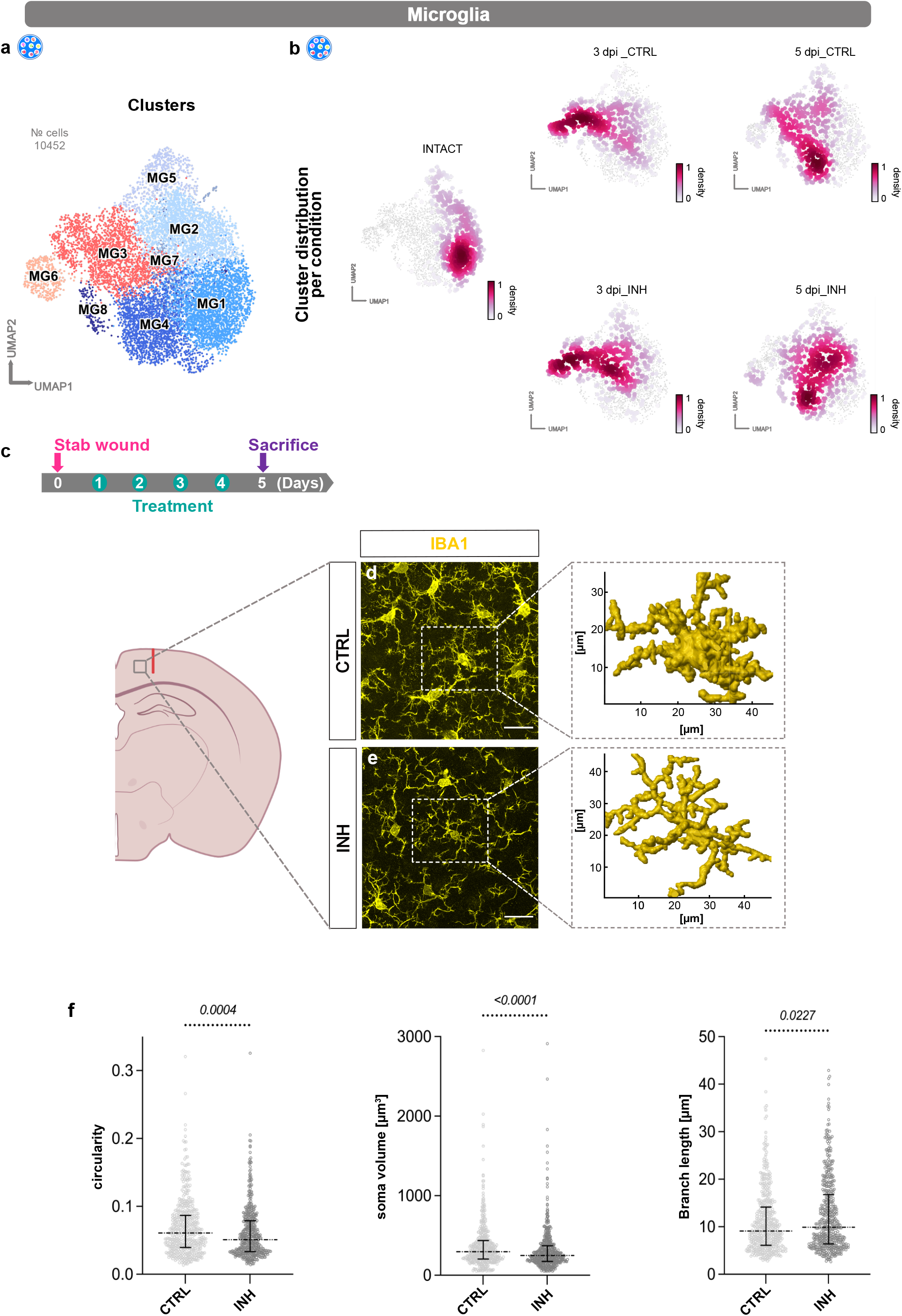
Cxcr3 and Tlr1/2 pathway inhibition after stab wound injury decreases microglial reactivity. (**a**,**b**) UMAPs illustrating subclusters of microglia (**a**) and cell distributions (**b**) among those subclusters at all time points (intact, 3 dpi, and 5 dpi) and conditions (CTRL and INH). Data are downsampled to an equal number of cells between time points and conditions (**b**). (**c**) Experimental paradigm for assessing microglial morphology characteristics according to Heindl et al.^54^. The gray box on mouse brain scheme highlights the analyzed area. The red line indicates the injury core. (**d**,**e**) Representative images of Iba1^+^ microglia (yellow) in CTRL (**d**) and INH-treated (**e**) mice. Dashed white boxes indicate selected microglia used for 3D reconstruction. All images are full z-projections of confocal z-stacks. (**f**) Scatter plot depicting morphological features of analyzed microglial cells. Each data point represents one microglial cell; a total of 450 cells per condition were analyzed. Data are displayed as median ± interquartile range. p-values were determined with Mann-Whitney U-test. Scale bars: **d**,**e**: 20 μm. Abbreviations: UMAP = uniform manifold approximation and projection, dpi = days post injury, CTRL = stab wound-injured control animals, INH = stab wound-injured inhibitor-treated animals.

In summary, our scRNA-seq data implied that Cxcr3 and Tlr1/2 pathway inhibition accelerates the transition from a reactive to a homeostatic microglial cell state early after injury. These findings were further supported by pronounced morphological changes in inhibitor-treated microglia, which are characteristic features of less reactive cells.

### Altered astrocyte response after Cxcr3 and Tlr1/2 pathway inhibition

To address the effects of Cxcr3 and Tlr1/2 pathway inhibition on astrocytes after brain injury, we subclustered astrocytes (Fig. 8a) and investigated the cell distribution among all conditions and time points (Fig. 8b). Astrocytes originating from intact conditions were evenly distributed among all homeostatic clusters. However, cells from stab-wounded brains were initially localized in both homeostatic and reactive clusters at 3 dpi, whereas at 5 dpi, most cells were distributed among the reactive clusters. Comparison of astrocyte cell distribution of SW CTRL and SW INH samples indicated noticeable differences at 5 dpi. Most cells originating from the SW CTRL condition were distributed among the reactive clusters AG5, AG6, and AG7, whereas cells originating from the SW INH condition were largely confined to the reactive cluster AG5 (Fig. 8b). Interestingly, cluster AG5 exhibited lower expression of reactivity markers, such as *Gfap* and *Lcn2*, than the reactive clusters AG6 and AG7 (E.D. Fig. 14b). In line with the shifted distribution of SW INH cells to cluster AG5, inhibitor-treated astrocytes also displayed lower expression of *Gfap* and *Lcn2* at 5 dpi (E.D. Fig. 14c). To determine whether astrocyte reactivity was altered overall, we generated astrocyte reactivity scores (based on Hasel et al.^36^) and compared the reactivity gene set scores among intact, stab wound-injured control, and inhibitor-treated samples (E.D. Fig. 14d). Generally, both reactivity scores (Cl4 and Cl8 in E.D. Fig 14d) were relatively lower in stab wound-injured inhibitor-treated samples at both time points (3 and 5 dpi). However, the fraction of astrocytes expressing these distinct gene sets was unchanged. Therefore, our analysis implied that the inhibitor treatment decreased astrocyte reactivity overall but was not sufficient to promote the return of reactive astrocytes to homeostasis. In line with our findings from the scRNA-seq analysis, also by immunohistochemical analysis, we did not observe differences in the total number of reactive astrocytes between stab-wounded control and inhibitor-treated mice at 5 dpi (E.D. Fig. 14e-i). Both experimental groups showed comparable GFAP^+^ cell accumulation (E.D. Fig. 14f, g, h), and numbers of NGAL^+^ (*Lcn2*) and GFAP^+^ positive astrocytes in the injury vicinity (E.D. Fig. 14f, g, i).

**Figure 8.**
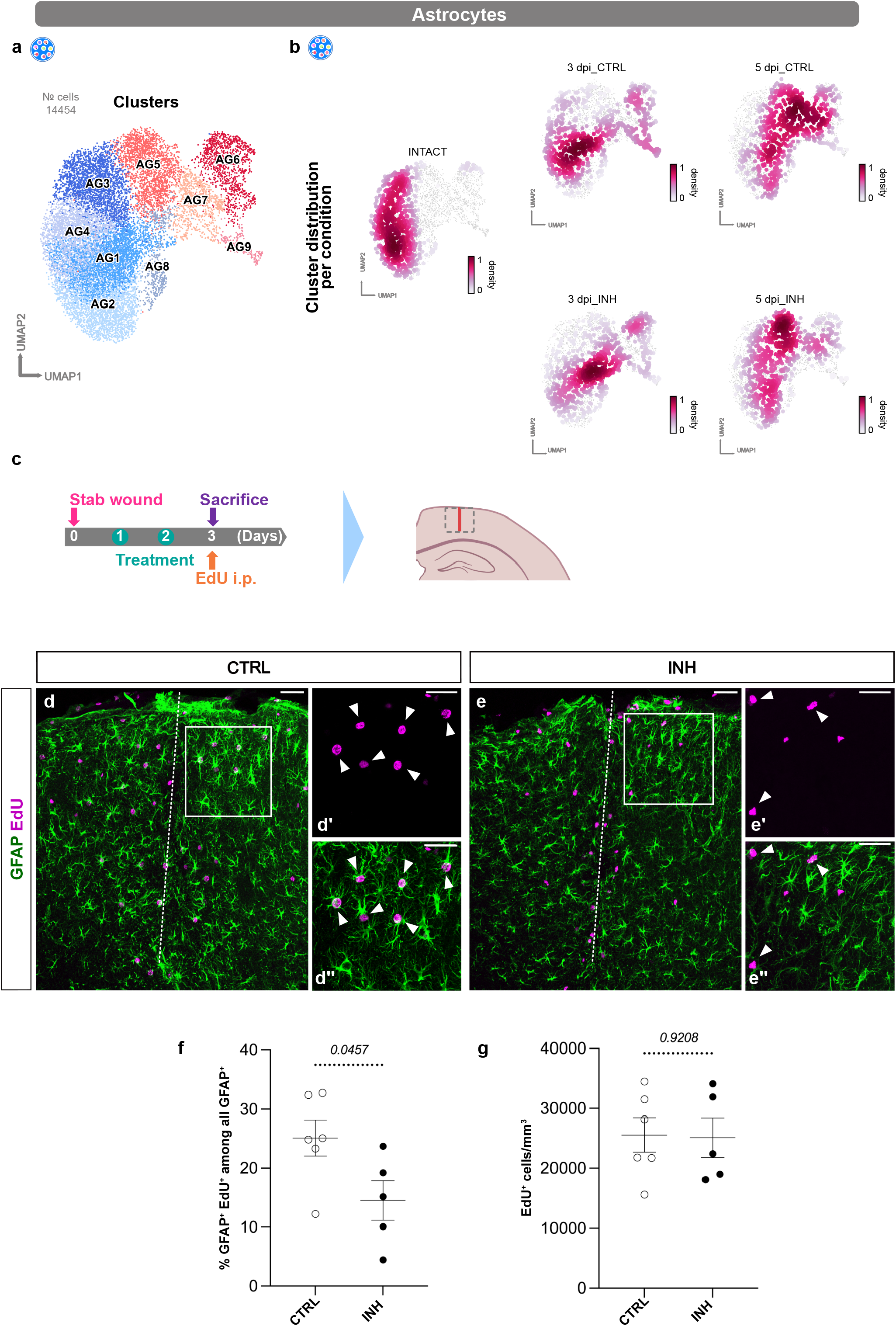
Proliferation of reactive astrocytes is decreased after Cxcr3 and Tlr1/2 pathway inhibition. (**a**,**b**) UMAPs illustrating subclusters of astrocytes (**a**) and cell distributions (**b**) among those subclusters at all time points (intact, 3 dpi, and 5 dpi) and conditions (CTRL and INH). Data are downsampled to an equal number of cells between time points and conditions (**b**). (**c**) Experimental paradigm for assessing astrocyte proliferation. The dashed gray box on mouse brain scheme indicates the analyzed area. The red line indicates the injury core. (**d**,**e**) Representative overview images of proliferating GFAP^+^ (green) and EdU^+^ (magenta) astrocytes in CTRL (**d**) and INH-treated (**e**) animals. White dashed lines highlight injury cores. Micrographs (**d’-e’’**) are magnifications of white boxed areas in (**d**) and (**e**), respectively. White arrowheads in micrographs indicate colocalization of EdU (**d’**,**e’**) with GFAP^+^ astrocytes (**d’’**,**e’’**). All images are full z-projections of confocal z-stacks. (**f**,**g**) Dot plots depicting percentage of proliferating (GFAP^+^ and EdU^+^) astrocytes (**f**) and total density of proliferating (EdU^+^) cells (**g**) in CTRL and INH-treated animals. Data are shown as mean ± standard error of the mean. Each data point represents one animal. p-values were determined with unpaired t-test. Scale bars: **d**,**e**: 50 μm (overview), **d’-e’’**: 20 μm (micrographs). Abbreviations: UMAP = uniform manifold approximation and projection, dpi = days post injury, EdU = 5-ethinyl-2′-deoxyuridine, i.p. = intraperitoneal injection, CTRL = stab wound-injured control animals, INH = stab wound-injured inhibitor-treated animals.

Furthermore, beyond the diminished expression of reactive astrocyte markers in cluster AG5, this cluster was also devoid of proliferating cells, because most cycling cells were confined to clusters AG6 and AG7, as indicated by the scRNA-seq proliferation score (E.D. Fig. 14j, Ext. Table 6). Interestingly, on the basis of our scRNA-seq analysis, interference with Cxcr3 and Tlr1/2 signaling after stab wound injury decreased the fraction of proliferating astrocytes at 3 and 5 dpi, in line with the abundance of SW INH cells composing cluster AG5 (Fig. 8b, E.D. Fig. 14k).

To further investigate potential alterations in proliferation after inhibitor treatment, we assessed astrocyte proliferation with immunohistochemistry in combination with the DNA-base analogue EdU (0.05 mg/g i.p. injection 1 hr before sacrifice) at 3 dpi (Fig. 8c). Indeed, inhibition of the Cxcr3 and Tlr1/2 pathways after injury significantly decreased the proportion of proliferating (GFAP^+^ and EdU^+^) astrocytes in the injury vicinity (Fig. 8d-f). However, the total number of EdU^+^ cells was not altered (Fig. 8d, e, g). In summary, our scRNA-seq analysis demonstrated decreased astrocyte reactivity and proliferation rates after inhibitor treatment. However, Cxcr3 and Tlr1/2 pathway inhibition, despite being sufficient to decrease astrocyte proliferation *in vivo*, did not completely revert astrocytes to homeostasis.

## Discussion

TBIs have complex pathophysiology involving responses of various types of cells^3,4^. However, most studies have focused on the responses of specific cell types, whereas few have evaluated the interplay among these cells^23,26,55^. Therefore, we developed a comprehensive dataset profiling the transcriptional changes across various cell types in spatial and temporal contexts. Our study used the stab wound injury model in mice^26,31^, a mild injury model involving breakdown of the BBB and the activation of both glial and immune cells^26^. Our model’s reproducibility and observed reactivity indicates its suitability for studying the basic features of TBI pathophysiology.

Spatial transcriptomic analysis of the stab-wounded cortex at 3 dpi revealed an injury-specific cluster, cluster VI, around the injury core without detectable changes in the cortex regions distant from the injury. The specificity of the injury defining the cluster VI signature was validated by the expression patterns of selected genes (*Serpina3n, Lcn2*, and *Cd68*) with RNAscope and immunohistochemistry, whose results were in line with those from the stRNA-seq analysis. This observation supports the use of stRNA-seq to detect global changes with spatial information. To complement the clustering analysis, we investigated gene expression patterns by conducting spatial gradient analysis. This analysis revealed gene expression changes in pre-defined gradients spanning from the injury core to the periphery, whereby injury-enriched genes followed various types of descending patterns. By analyzing the gene sets following these descending patterns, we observed an over-representation of processes associated with innate immunity. Complementing these results, cluster VI-enriched genes also revealed the regulation of processes associated with the immune system, in addition to angiogenesis and phagocytosis. Clearing dead cells and debris and re-establishing vasculature to ensure sufficient oxygen supply are critical defense mechanisms that occur early after brain damage^32^.

Many observed local changes represented by the injury-enriched genes were associated with reactive astrocytes and microglia^12,34–37^, thus indicating an overrepresentation of these populations in the injury milieu. Interestingly, we did not identify reactive OPC hallmarks despite clear evidence of reactive OPCs at the injury site^8,56^. This result may be partly explained by the unknown signature of reactive OPCs, because only an increase in proliferation and expression of CSPG4 have been used to identify reactive OPCs to date^9^. Moreover, the combination of stRNA-seq with scRNA-seq analysis is becoming an excellent tool to reveal transcriptomic changes in specific cell types in relation to their predicted location. This capability is of great interest for any focal pathology, given that the reactions of astrocytes^16,^ OPCs^11,57^, and microglia^57,58^ have been found to depend on their distance from the pathological site. Integration of scRNA-seq and stRNA-seq datasets indicated the presence of distinct cell types in the injury environment. We detected multiple populations responding to the injury by enriched or decreased representation in the injured milieu, whereas other cell clusters never responded. Microglial clusters displayed a uniform response to injury because all microglia clusters were found to accumulate at the injury site, and cluster 11_Microglia had the highest correlation. The activation of microglia was consistent with our immunohistochemical analysis findings but differed from the specific activation patterns observed in the APP model of neurodegeneration^57^. In the APP model, certain cells display elevated expression of the disease-associated microglia signature concentrated in areas of plaque deposition, as determined by stRNA-seq. Contrary to microglia, astrocytes showed a heterogeneous response, with clusters 12_Astrocytes and 23_Astrocytes responding to injury and accumulating around the injury site. In contrast, the remaining astrocytic clusters were underrepresented in the injury area with respect to the rest of the cortex. Interestingly, the injury-enriched astrocytic clusters 12_Astrocytes and 23_Astrocytes displayed unique features corresponding to their location and gene signatures. Cluster 12_Astrocytes for example, expressed high levels of *Gfap*, whereas cluster 23_Astrocytes might represent the recently described atypical astrocytes, which, after focal brain injury, rapidly downregulate GFAP and other astrocytic proteins^30^. Generally, OPCs also responded to the injury, however, cluster 15_OPCs was the only cluster showing enrichment at the injury core. Finally, we detected the responses of peripheral infiltrating macrophages and monocytes and found that clusters 13_Macrophages/Monocytes and 18_Monocytes contributed to the injury milieu. With the integration of the two datasets, we were able to identify the cells populating the injury core, thus offering a possibility for further thorough investigations.

The addition of the dataset generated at 5 dpi allowed us to analyze the temporal changes in response to injury. Microglia displayed elevated reactivity at 3 dpi, whereas at 5 dpi, cells shifted toward the homeostatic clusters. OPCs were characterized by a fast transient response at 3 dpi as very few cells resided in the reactive cluster at 5 dpi. In contrast, astrocyte reactivity peaked at 5 dpi, as most cells at this time point were confined to the reactive clusters, whereas at 3 dpi, many astrocytes still resided in the homeostatic clusters. Together, our data suggest that the activation of glial cells is continuous and does not involve the appearance or disappearance of distinct clusters at any specific time point. Therefore, our resource provides an excellent opportunity to investigate the processes driving cellular reactivity in response to injury in detail.

To provide a proof of principle, we comprehensively examined the genes characterizing each subcluster. In this way, we identified subclusters of astrocytes, microglia and OPCs with shared enriched signatures, including proliferation and activation of innate immune processes. Indeed, proliferation is a hallmark of injury-induced reactivity, including microglia^59^, astrocytes^34,60^, and OPCs^8,61^, thus further validating our approach. The shared inflammatory signature identified in this study is largely present in other brain pathologies, such as in LPS-induced reactive astrocytes^36^. However, although a high proportion of shared inflammatory genes were expressed in the reactive astrocyte cluster 8, not all shared inflammatory genes were detected. This finding implies a common expression of core innate immunity-associated genes in different cell types in response to a variety of stimuli (e.g., LPS or TBI). However, each pathological condition further triggers distinct inflammatory-associated processes, which are uniquely coordinated in each pathology. The inflammatory signature present in the reactive clusters MG4, AG5, and OPCs2 included several IFN-I pathway genes. Among these genes, Interferon regulatory factor 7 (*Irf7*), a transcription factor crucial for IFN-I activity^62^, and *Cxcl10*, a well-characterized ligand of the Cxcr3 pathway^47^, were detected. Previous studies have demonstrated that Irf7 induces type I IFNs through the activation of TLR2, thus resulting in the transcription of several mediators, including *Cxcl10*^63,64^. In addition, the Tlr2/Irf7 signaling axis has been associated with microglia-mediated inflammation after subarachnoid hemorrhage in mice^65^. Furthermore, we have recently demonstrated that Cxcr3 and Tlr1/2 regulate OPC accumulation at injury site in the zebrafish brain in a redundant and synergistic manner^49^. Our data support the novel concept that the same innate immune pathways are responsible for initiating the response in injury-induced reactive glial clusters.

Because both pathways were regulated in several reactive populations, we systemically inhibited the above-mentioned pathways after brain injury by treating the animals simultaneously with a specific antagonist for Cxcr3^50^ (NBI-74330) and a Tlr1/2 pathway inhibitor (CU CPT 22)^51^. We then performed scRNA-seq analysis at 3 and 5 dpi and integrated the datasets with our control analysis. This allowed us to investigate cell type-specific changes emerging after inhibitor treatment. Interestingly, we observed that multiple innate immunity genes, including *Irf7*, were downregulated after inhibition of the Cxcr3 and Tlr1/2 pathways, and clusters AG7, MG3, and OPCs2 were affected the most. These results are in line with our observation that up-regulation of these genes is associated with the emergence of these clusters after injury. However, whether the downregulation of the shared inflammation signature is a direct consequence of the Tlr1/2 and Cxcr3 inhibition in cells themselves, or a consequence of altered cell-cell communication after systemic Tlr1/2 and Cxcr3 inhibition, remains unclear. Additionally, by performing differential gene expression analysis within each subcluster between SW CTRL and SW INH conditions at each time point, we addressed changes induced in each cluster separately. Notably, in response to the inhibitor treatment, innate immune processes were generally downregulated in astrocytes, microglia, and OPCs at 3 dpi, whereas at 5 dpi, the regulation became specific to each cell population. This finding suggests that a shared initial regulation diverges and consequently drives specific reactions in each cell type. In conclusion, we demonstrated that inhibition of the Cxcr3 and Tlr1/2 pathways modulates innate immunity in glial cells on a temporal basis.

Next, we examined how the systemic inhibition of the two pathways affected the reactivity of glial cells. Thus, we addressed the numbers of oligodendrocytes (OLIG2^+^ cells) together with their proliferation. Surprisingly, we did not observe any difference between SW CTRL and SW INH conditions at 3 dpi. This observation contrasts with the findings from our study in zebrafish, in which inhibition of both pathways was found to alleviate reactive gliosis by decreasing the accumulation of oligodendrocytes and their proliferation^49^. This lack of concordance might be due to differences in the injury environment, and additional pathways involved in the accumulation and proliferation of OPCs in the mouse cerebral cortex and may reflect a possible difference between regeneration competent and regeneration incompetent species.

In contrast, in examining microglial reactivity via morphological analysis^54^, we observed that inhibitor-treated microglia were in a less reactive/activated state than stab wound-injured controls. Specifically, microglia originating from SW INH animals displayed significantly smaller cell somata, a less round shape, and increased branch length, and appeared to be more ramified than microglia of SW CTRL animals. This observation, in combination with the cell distribution in the scRNA-seq analysis, suggests that blocking the Tlr1/2 and Cxcr3 pathways accelerates the microglial transition from a reactive to a homeostatic state.

Similarly, astrocytes showed altered reactivity when the Cxcr3 and Tlr1/2 signaling pathways were inhibited: inhibitor-treated astrocytes displayed a decrease in expression of reactivity markers. Additionally, we observed a decrease in the number of proliferating astrocytes in the injury vicinity at 3 dpi. However, at 5 dpi, we did not observe differences in the overall astrocyte reactivity state between SW CTRL and SW INH, as indicated by our transcriptomic analysis and the follow-up immunohistochemical assessments. These findings suggested a coordinated response of astrocytes, and presumably glial cells in general, whereby different pathways regulate distinct aspects of glial reactivity. Individual signaling pathways might potentially even be involved in highly divergent functions in different glial cells. In such a scenario, the Tlr1/2 and Cxcr3 pathways may largely regulate proliferation in astrocytes, and cellular morphological aspects in microglia, in line with different brain pathologies inducing various glial responses (for example astrocyte proliferation)^60,66,67^. These findings further emphasize the versatility of our datasets and analysis.

In conclusion, the present study provides a comprehensive transcriptomic dataset for analyzing early events after TBI with respect to changes in time, space, and cell type. Additionally, this dataset provides an excellent platform to examine the interplay of a variety of cells in response to injury. A better understanding of injury pathophysiology may provide more opportunities for developing new therapeutic strategies.

## Acknowledgments

We are particularly grateful to the entire Neurogenesis and Regeneration group members for their experimental inputs and discussions and Dr. Alessandro Zambusi for critical reading of the manuscript. We acknowledge the support of the following core facilities: the Bioimaging Core Facility and Bioinformatic Core Facility at the BioMedical Center of LMU Munich, the Laboratory for Functional Genome Analysis (LAFUGA), and the Sequencing Facility at the Helmholtz Zentrum München. This work was supported by the German research foundation (DFG) by the SFB 870; TRR274; SPP 1738 “Emerging roles of non-coding RNAs in nervous system development, plasticity & disease”, SPP1757 “Glial heterogeneity”; SPP2191 “Molecular mechanisms of functional phase separation” (ID 402723784) and the Excellence Strategy within the framework of the Munich Cluster for Systems Neurology (EXC 2145/1010 SyNergy – ID 390857198), Fritz Thyssen Stiftung and Ampro Helmholtz Alliance.

## Author Contributions

C.K., V.Z. and J.N. conceived the project and experiments. C.K., V.Z., J.F.S., T.S.E. and R.B. performed experiments and analyzed data. C.K., H.A. and S.F. performed the bioinformatic analyses C.K., V.Z. and J.N. wrote the manuscript with input from all authors. J.N., M.G., M.D. and F.J.T. supervised research and acquired funding.

## Figure Legends

**E.D. Figure 1. Spatial transcriptome of intact and gray matter stab wound-injured mice**.

(**a-b**) Spatial transcriptomics in intact and stab wound-injured mice (3 dpi). Brains were manually resected and positioned on 10x Visium capture areas. In each capture area, two brain sections of either intact (**a**) or stab wound-injured cortices (**b**) were collected. (**c**) Clustering of gene expression data on spatial coordinates based on highly variable genes and subsequent dimensionality reduction. (**c**) Dot plot depicting the 5 most enriched genes in each of the 16 identified clusters. (**e**) Expression of neuronal layer scores 2/3, 4, 5 and 6 in intact and injured brain sections on spatial coordinates. The white dashed lines highlight cluster VI borders. Neuronal layer gene set scores are listed in Ext. Table 2. (**e**) Heatmap depicting the expression of the 25 most ascending genes along the spatial trajectory (see Fig. 2) depicted as mean expression of injury 1 and 2. (**f**) Heatmap displaying the 30 most enriched ascending gene sets along the spatial trajectory (see Fig. 2) depicted as mean expression from injury 1 and 2. Abbreviations: BP = biological processes, MF = molecular functions, CC = cellular components, GO = gene ontology, REACT = reactome, dpi = days post injury.

**E.D. Figure 2. scRNA-seq clustering of intact and brain-injured mice (3 dpi) and cell-type identification**.

(**a**) UMAP plots depicting clustering of cells originating from intact (2676 cells) and injured (3646 cells) conditions among 30 defined clusters. Clusters are color-coded and annotated according to their transcriptional identities. Note that clusters 13_Macrophages/Monocytes, 17_DCs, and 23_Astrocytes are absent in the intact condition. (**b**) Dot plot depicting the expression of the 3 most enriched genes in each of the 30 identified clusters (see Ext. Table 5). (**c-e**) UMAPs highlighting example marker genes to identify microglia/macrophages (**c**), astrocytes (**d**) OPCs (**e**) and cycling cells (**f**). G2/M gene set score is listed in Ext. Table 6. Abbreviations: UMAP = uniform manifold approximation and projection, dpi = days post injury, OPCs = oligodendrocyte progenitor cells, COPs = committed oligodendrocyte progenitors, MOL = mature oligodendrocytes, VECV = vascular endothelial cells (venous), VSMCs = vascular smooth muscle cells, VLMCs = vascular and leptomeningeal cells, BAM = border-associated macrophages, NKT cells = natural killer T cells, DCs = dendritic cells.

**E.D. Figure 3. Probabilistic mapping of in scRNA-seq identified cellular clusters on Visium dataset**.

(**a**) Probabilistic mapping of in Fig. 2b identified scRNA-seq clusters on the spatial transcriptome dataset (3 dpi). Stab wound injury elicits different mode of reaction in the injury vicinity. Plots are grouped into injury enriched clusters (upper panel) and remaining clusters (lower panel). Abbreviations: OPCs = oligodendrocyte progenitor cells, NKT cells = natural killer T cells, DCs = dendritic cells, MOL = mature oligodendrocytes, VSMCs = vascular smooth muscle cells, COPs = committed oligodendrocyte progenitors, BAM = border-associated macrophages, VLMCs = vascular and leptomeningeal cells, VECV = vascular endothelial cells (venous).

**E.D. Figure 4. Single-cell deconvolution-based mapping of in scRNA-seq identified cellular clusters on Visium dataset**.

(**a**,**b**) Deconvolution based mapping of in E.D. Fig. 3 identified scRNA-seq clusters on the spatial transcriptome dataset (3 dpi). Plots are grouped into injury enriched clusters (upper panel) and remaining clusters (lower panel) (**b**). Abbreviations: OPCs = oligodendrocyte progenitor cells, NKT cells = natural killer T cells, DCs = dendritic cells, MOL = mature oligodendrocytes, VSMCs = vascular smooth muscle cells, COPs = committed oligodendrocyte progenitors, BAM = border-associated macrophages, VLMCs = vascular and leptomeningeal cells, VECV = vascular endothelial cells (venous).

**E.D. Figure 5. Integration of scRNA-seq datasets of intact and brain-injured mice (3 and 5 dpi), cluster distribution and cell type identification**.

(**a**,**b**) UMAP plots depicting clustering (**a**) and cell distribution of intact (16567 cells), 3 dpi (3637 cells), and 5 dpi (13658 cells) cells among 35 defined clusters (**b**). Clusters were color-coded and annotated according to their transcriptional identities. (**c**) Dot plot depicting expression of the 3 most enriched genes in each of the 35 identified clusters (see Ext. Table 8). Abbreviations: UMAP = uniform manifold approximation and projection, dpi = days post injury, VECV = vascular endothelial cells (venous), MOL = mature oligodendrocytes, DAM = disease-associated microglia, OPCs = oligodendrocyte progenitor cells, NKT cells = natural killer T cells, BAM = border-associated macrophages, VSMCs = vascular smooth muscle cells, COPs = committed oligodendrocyte progenitors, VLMCs = vascular and leptomeningeal cells, DC = dendritic cells.

**E.D. Figure 6. Stab wound injury elicits unique gene expression in distinct glial subpopulations**.

(**a**) UMAP depicting subclustered astrocytes of integrated and batch-corrected intact, 3 dpi, and 5 dpi datasets. (**b**) Dot plot depicting expression of the 5 most enriched genes in each of the 9 identified clusters. (**c**) UMAP displaying cell distribution of intact (green), 3 dpi (red) and 5 dpi (orange) cells among all 9 astrocytic clusters. Note that clusters AG5, AG6, and AG8 are mainly composed of cells originating from injured cortices. (**d**) UMAP displaying localization of injury-enriched cluster 12_Astrocytes (turquoise, E.D. Fig. 4b) on subclustered astrocytes. (**e**) UMAPs highlighting expression of example marker genes *Gfap, Vim* and *Lcn2* to identify reactive astrocyte clusters. (**f**) UMAP depicting subclustered microglia of integrated and batch-corrected intact, 3 dpi, and 5 dpi datasets. (**g**) Dot plot depicting expression of the 5 most enriched genes in each of the 8 identified clusters. (**h**) UMAP displaying cell distribution of intact (green), 3 dpi (red), and 5 dpi (orange) cells among all 8 microglial clusters. Note that clusters MG4 and MG6 are mainly composed of cells originating from injured cortices. (**i**) UMAP displaying localization of injury-enriched cluster 11_Microglia (peach, E.D. Fig. 4b) on subclustered microglia. (**j**) UMAPs highlighting expression of example marker genes *Aif1, Tmem119* and *P2ry12* to identify reactive microglial clusters. (**k**) UMAP depicting subclustered oligodendrocytes of integrated and batch-corrected intact, 3 dpi, and 5 dpi datasets. (**l**) Dot plot depicting expression of the 5 most enriched genes in each of the 10 identified clusters. (**m**) UMAP displaying cell distribution of intact (green), 3 dpi (red), and 5 dpi (orange) cells among all 10 clusters. Note that cluster OPCs2 is mainly composed of cells originating from both injured conditions. (**n**) UMAP displaying localization of injury-enriched cluster 15_OPCs (pink, E.D. Fig. 4b) on subclustered oligodendrocytes. (**o**) Dot plot depicting expression of the 30 most enriched genes in cluster OPCs2. Note comparable gene expression of clusters OPCs1 and OPCs2 prevent the identification of unique reactive OPC markers. Abbreviations: UMAP = uniform manifold approximation and projection, dpi = days post injury, OPCs = oligodendrocyte progenitor cells, COPs = committed oligodendrocyte progenitors, MFOL = myelin-forming oligodendrocytes, MOL = mature oligodendrocytes.

**E.D. Figure 7. Reactive glial subpopulations share injury-induced transcriptomic signature**.

(**a**,**b**) UpSet plots depicting unique and overlapping downregulated (**a**) and upregulated DEGs (**b**) (pval < 0.05, log_2_ fold change > 1.6 or log_2_ fold change < -1.6) between different glial subclusters of integrated and batch-corrected intact, 3 dpi, and 5 dpi datasets (Fig. 5c-e). Wherever applicable, DEGs are determined by comparing each glial subcluster to all other subclusters within the respective cell type (further details in Methods). The red bars in (**b**) highlight commonly shared genes between reactive astrocytic, microglial, and oligodendroglial subclusters. Abbreviations: DEGs = differentially expressed genes.

**E.D. Figure 8. Reactive glial subpopulations display shared gene expression following injury**.

(**a**) Experimental paradigm for assessing shared gene expression in astrocytes, microglia (*C57Bl6/J*), and OPCs (*NG2-CreER*^*T2*^*xCAG-GFP*). The dashed gray box on mouse brain scheme indicates the example imaging area. The red line indicates the injury core. (**b-g**) RNAscope example images of shared innate immunity-associated genes *Oasl2* (**b-d**) and *Ifi27l2a* (**e-g**) (magenta) counterstained with GFAP (black) (**b**,**e**), IBA1 (black) (**c, f**), and GFP (NG2^+^ glia) (black) (**d**,**g**) antibodies. Micrographs (**b’-g’’**) are magnifications of the red boxed areas in corresponding overview images. The orange arrowheads in micrograph depict double positive cells. All images are single z-plane of confocal z-stacks. Scale bars: (**e-g”**): 20 μm.

**E.D. Figure 9. Reactive glial subpopulations display shared marker expression after injury**.

(**a**) Experimental paradigm for assessing shared marker expression in astrocytes, OPCs, and microglia. The dashed gray box on mouse brain scheme indicates the analyzed area. The red line illustrates the injury core. (**b-d**) Representative overview images depicting Galectin1 expression in GFAP^+^ astrocytes (**b**), NG2^+^ glia (GFP) (**c**), and IBA1^+^ microglia (**d**). Micrographs (**b’-d’’**) are magnifications of white boxed areas in corresponding overview images. The white dashed lines indicate injury cores. The white arrowheads in the micrographs depict colocalization of Galactin1 (**b’-d’**) with GFAP^+^ (**b’’**), GFP^+^ (**c’’**), and IBA1^+^ (**d’’**) cells. All images are full z-projections of confocal z-stacks. (**e**) Heatmap depicting the shared inflammatory signature gene score between conditions and clusters. (**f**) UMAPs depicting localization of the shared inflammatory signature gene scores among all defined cell clusters. (**g**) Dot plot depicting gene expression of components associated with the Tlr1/2 and Cxcr3 signaling pathways. Genes are adapted and complemented based on Sanchez-Gonzalez et al.^49^.Scale bars: **b-d**: 50 μm (overview), **b’-d’’**: 20 μm (micrographs).

**E.D. Figure 10. Common inflammatory gene signature in murine LPS- and human iPSC-induced reactive astrocytes**

(**a**,**b**) UMAPs illustrating selective marker gene expression of shared inflammatory signature (see Fig. 5c) in subclustered astrocytes based on Hasel et al. 2021. Plots depicting presence (**a**) and absence (**b**) of several shared inflammatory genes in reactive astrocyte cluster 8. (**c**,**d**) UMAPs depicting localization of IRAS1 and IRAS2 gene scores (**c**) among all defined cell clusters (**d**). IRAS1 and IRAS2 gene scores are based on Leng et al.^48^. Abbreviations: UMAP = uniform manifold approximation and projection, CNT = control, LPS = lipopolysaccharide, Exp = expression, SW = stab wound, IRAS = inflammatory reactive astrocytes signature.

**E.D. Figure 11. scRNA-seq data integration of intact and stab wound-injured cortices**.

(**a**) UMAP plots depicting cell distribution of integrated and batch-corrected intact (green, 16649 cells), 3 dpi CTRL (red, 3637 cells), 3 dpi INH (pink, 4613 cells), 5 dpi CTRL (orange, 13766 cells), and 5 dpi INH (peach, 15146 cells) datasets. (**b**) Heatmap displaying high cluster similarity of integrated intact and injured CTRL conditions (control, y-axis), and integrated intact, injured CTRL, and injured INH conditions (control and inhibitor-treatment, x-axis). (**c**) UMAP displaying cell distribution of intact (green), 3 dpi CTRL (red), 3 dpi INH (pink), 5 dpi CTRL (orange), and 5 dpi INH (peach) cells among the 9 identified astrocytic clusters as depicted in Fig. 5c. Clusters AG5, AG6, AG7 and AG8 are mainly formed by cells originating from injured CTRL and INH animals. (**d**) UMAP displaying cell distribution of intact (green), 3 dpi CTRL (red), 3 dpi INH (pink), 5 dpi CTRL (orange), and 5 dpi INH (peach) cells among the 8 identified microglial clusters as depicted in Fig. 5d. Clusters MG3 and MG6 are mainly formed by cells originating from injured CTRL and INH animals. (**e**) UMAP displaying cell distribution of intact (green), 3 dpi CTRL (red), 3 dpi INH (pink), 5 dpi CTRL (orange), and 5 dpi INH (peach) cells among the 13 identified oligodendroglial clusters as depicted in Fig. 5e. Clusters OPCs1 and OPCs2 are mainly formed by cells originating from injured CTRL and INH animals. Abbreviations: UMAP = uniform manifold approximation and projection, dpi = days post injury, CTRL = stab wound-injured control mice, INH = stab wound-injured inhibitor-treated mice, VECV = vascular endothelial cells (venous), MOL = mature oligodendrocytes, OPCs = oligodendrocyte progenitor cells, NKT cells = natural killer T cells, BAM = border-associated macrophages, VSMCs = vascular smooth muscle cells, COPs = committed oligodendrocyte progenitors, VLMCs = vascular and leptomeningeal cells, DC = dendritic cells.

**E.D. Figure 12. The Cxcr3 and Tlr1/2 pathway inhibition after brain injury induces subcluster specific changes**.

(**a-d**) UpSet plots depicting unique and overlapping downregulated (**a**,**c**) and upregulated (**b**,**d**) DEGs (pval < 0.05, log_2_ fold change > 0.7 or log_2_ fold change < - 0.7) between different glial subclusters in response to the Cxcr3 and Tlr1/2 inhibition at 3 dpi (**a**,**b**) and 5 dpi (**c**,**d**). (**e**,**f**) GO term networks illustrate distinct, subcluster-specific biological processes enriched in the set of genes upregulated in response to the Cxcr3 and Tlr1/2 pathway inhibition at 3 dpi (**e**) and 5 dpi (**f**) (see Ext. Table 12). Abbreviations: DEGs = differentially expressed genes, dpi = days post injury.

**E.D. Figure 13. OPC reactivity after injury is not altered by the Cxcr3 and Tlr1/2 pathway inhibition**.

(**a**,**b**) UMAPs illustrating subclusters of oligodendrocytes (**a**) and cell distributions among these subclusters (**b**) at all time points (intact, 3 dpi, and 5 dpi) and conditions (CTRL and INH). Data are downsampled to an equal number of cells between time points and conditions (**b**). (**c**) Experimental paradigm for assessing number of oligodendrocytes and OPC proliferation in injury vicinity. The dashed gray box on mouse brain scheme indicates the analyzed area. The red line highlights the injury core. (**d**,**e**) Representative overview images of proliferating OLIG2^+^ (gray) and EdU^+^ (magenta) oligodendrocytes in CTRL (**d**), and INH-treated (**e**) animals. The white dashed lines highlight injury cores. Micrographs (**d’-e’’**) are magnifications of the white boxed areas in (**d**) and (**e**), respectively. The white arrowheads in micrographs depict colocalization of EdU (**d’**,**e’**) with OLIG2^+^ (**d’’**,**e’’**) cells. All images are full z-projections of confocal z-stacks. (**f**,**g**) Dot plots depicting number of oligodendrocytes (OLIG2^+^ cells) (**f**) and proliferating OPCs (OLIG2^+^ and EdU^+^) (**g**) in CTRL and INH-treated animals. Data are shown as mean ± standard error of the mean. Each data point represents one animal. p-values were determined with unpaired t-test. Scale bars: **d**,**e**: 50 μm (overview), **d’**,**e’’**: 20 μm (micrographs). Abbreviations: UMAP = uniform manifold approximation and projection, dpi = days post injury, EdU = 5-Ethinyl-2′-deoxyuridine, i.p. = intraperitoneal injection, CTRL = stab wound-injured control animals, INH = stab wound-injured inhibitor-treated animals, OPCs = oligodendrocyte progenitor cells, COPs = committed oligodendrocyte progenitors, MFOL = myelin-forming oligodendrocytes, MOL = mature oligodendrocytes.

**E.D. Figure 14. Reaction of microglia and astrocytes to stab wound injury in absence of the Cxcr3 and Tlr1/2 signaling**.

(**a**) Scatter plots depicting morphological microglial features. Each data point represents one microglial cell and in total 450 cells per condition were analyzed. Data are displayed as median ± interquartile range. p-values were determined with Mann-Whitney U-test. (**b**) UMAPs highlighting expression of *Gfap* and *Lcn2* among all astrocytic subclusters. (**c**) Dot plots depicting expression levels of *Gfap* and *Lcn2* in astrocytes. (**d**) Dot plot depicting astrocyte reactivity scores between all time points (intact, 3 dpi, and 5 dpi) and conditions (CTRL and INH). Genes defining astrocyte reactivity scores in E.D. Fig. 12d were extracted from cluster_4 and cluster_8 astrocytes of Hasel et al. 2021^36^ (Top 20 genes) and are plotted only in the reactive clusters (AG5, AG6, AG7 and AG9) (**e**) Experimental paradigm for assessing astrocyte reactivity in the injury vicinity. The dashed gray box on mouse brain scheme indicates the analyzed area. The red line highlights the injury core. (**f**,**g**) Representative overview images of GFAP^+^ astrocytes (green) and NGAL^+^ cells (magenta) in CTRL (**f**) and INH-treated (**g**) animals. White dashed lines highlight injury cores. Micrographs (**f’-g’’**) are magnifications of white boxed areas in (**f**) and (**g**), respectively. The white arrowheads in micrographs depict colocalization of NGAL (**f’**,**g’**) with GFAP^+^ astrocytes (**f’’**,**g’’**). All images are full z-projections of confocal z-stacks. (**h**,**i**) Dot plots depicting percentage of area covered with GFAP^+^ signal (**h**) and density of GFAP^+^ NGAL^+^ double positive astrocytes (**i**) in injury vicinity of CTRL and INH-treated mice. Data are shown as mean ± standard error of the mean. Each data point represents one animal. p-values were determined with unpaired t-test. (**j**) UMAPs highlighting localization of proliferating astrocytes (pink) to subclusters AG6 and AG7. (**k**) Histogram illustrating percentage of proliferating astrocytes between all time points (intact, 3 dpi, and 5 dpi) and condition (CTRL and INH). Proliferating astrocytes were identified in scRNA-seq datasets by S+G2/M score expression (see Ext. Table 6). Scale bars: **f**,**g**: 50 μm (overview), **f’-g’’**: 20 μm (micrographs). Abbreviations: UMAP = uniform manifold approximation and projection, dpi = days post injury, INT = intact mice, CTRL = stab wound-injured control animals, INH = stab wound-injured inhibitor-treated animals.

## Materials and Methods

### Animals

All operations were performed on 8-12 weeks old *C57Bl6/J* male mice, housed, and handled under the German and European guidelines for the use of animals for research purposes. Experiments were approved by the institutional animal care committee and the government of Upper Bavaria (ROB-55.2-2532.Vet_02-20-158). Anesthetized animals received a stab wound lesion in the cerebral cortex as previously described^31^, by inserting a thin knife into the cortical parenchyma using the following coordinates from Bregma: RC: -1.2; ML: 1-1.2 and from Dura: DV: -0.6 mm. To produce stab lesions, the knife was moved over 1mm back and forth along the anteroposterior axis from -1.2 to -2.2 mm. Animals were sacrificed 3 and 5 days after the injury (dpi).

For the treatment experiments, animals received inhibitors by gavage feeding. NBI 74330 (100 mg/kg, R&D Systems #4528) and CU CPT 22 (3 mg/kg, R&D Systems #4884) were dissolved in DMSO and diluted in corn oil. The vehicle solution consisted of DMSO diluted in corn oil and was administered to all control animals. To analyze the proliferative capacity of glial cells we injected 5-Ethinyl-2′-deoxyuridine (EdU, 0.05 mg/g, Thermofisher #E10187) intraperitoneally and animals were sacrificed 1hr after injection.

For the induction of Cre-mediated recombination in *NG2-CreER*^*T2*^*xCAG-GFP* mice, tamoxifen (40 mg/ml, Sigma #T5648) was administered orally. Animals received tamoxifen every second day (400 mg/kg) for a total of 3 times. Mice were injured two weeks after the last tamoxifen administration and sacrificed at 3dpi.

### Tissue preparation

Mice were deeply anesthetized and transcardially perfused with phosphate-buffered saline (PBS) followed by 4% paraformaldehyde (PFA) (wt/vol) dissolved in PBS. Brains were postfixed in 4% PFA overnight at 4°C, washed with PBS and cryoprotected in 30% sucrose at 4°C. Mouse brains used to assess microglia morphology were embedded in 3% agarose and cut coronally at 100 μm thickness using a vibratome (HM 650V, Microm). Otherwise, brains were embedded in frozen section medium Neg-50 (Epredia #6502), frozen and subsequently sectioned using a cryostat (Thermo Scientific CryoStar NX50). Coronal sections were collected either at a thickness of 20μm on slides for RNAscope or 40μm for free-floating immunohistochemistry.

### Immunohistochemistry

For immunohistochemistry, sections were blocked and permeabilized with 10% normal goat serum (NGS, vol/vol, Biozol, S-1000)/donkey serum (NDS, vol/vol, Sigma Aldrich 566460) and 0.5% Triton X-100 (vol/vol), dissolved in 1xPBS while being incubated overnight at 4°C with the corresponding primary antibodies. Following primary antibodies were used: anti-CD68 (rat 1:600, BioRad, MCA1957T), anti-Galactin1 (rabbit 1:200, Abcam, ab138513), anti-GFP (chick 1:400, Aves Labs, GFP-1020), anti-GFAP (goat 1:300, Abcam, ab53554), anti-GFAP (mouse 1:500, Sigma, G3893), anti-Iba1 (rabbit 1:500, Wako, 019-19741), anti-NGAL (rabbit 1:500, Thermofisher, PA5-79590), anti-SerpinA3n (goat 1:500, R&D Systems AF4709-SP). Sections were washed with PBS and incubated with secondary antibodies dissolved in 1xPBS solution containing 0.5% Triton X for two hours at room temperature. Following secondary antibodies were used: donkey anti-chick IgY A488 (1:1000, Dianova 703-545-155), goat anti-mouse IgG1 A546 (1:1000, Thermofisher A-21123), goat anti-rabbit IgG A546 (1:1000, Thermofisher A-11010), goat anti-rabbit IgG A633 (1:1000, Thermofisher A-21070), goat anti-rat IgG A488 (1:1000, Thermofisher A-11006). For nuclear labelling, sections were incubated with DAPI (final concentration of 4 μg/mL, Sigma, D9542) for 10 min at room temperature. EdU incorporation was detected by Click-iT™ EdU Alexa Fluor™ 647 Imaging Kit (Thermo Fisher Scientific #C10340) according to the manufacturer’s instructions. Staining procedure for microglia morphology analysis was performed as described in Heindl et al.^54^. Stained sections were mounted on glass slides with Aqua-Poly/Mount (Polysciences #18606).

### In situ hybridization

RNA in situ hybridization was performed using RNAscope® Multiplex Fluorescent Reagent Kit (ACD) according to the manufacturer’s instructions. Briefly, brain sections were fixed in 4% paraformaldehyde at 4 °C for 15 min, ethanol-dehydrated, treated with H_2_O_2_ and protease-permeabilized for 20min at 40 °C. Brain sections were then incubated for 2 h at 40 °C using the following probes: *Ifi27l2a*: 88617, *Serpina3n*: 430191-C2, *Lcn2:* 313971-C3, *Cd68:* 316611-C2, *Oasl2:* 534501, *Cxcl10*: 408921-C3. Signal was amplified according to the manufacturer’s instructions (Cat.Nr: 320293). Subsequently, sections were processed with immunohistochemistry analysis as described above. The primary antibodies used in combination with RNAscope® were as follows: chick antibody to GFP (1:500, Aves Lab, GFP-1020), goat antibody to GFAP (1:300, Abcam, ab53554), rabbit antibody to Iba1 (1:500, Wako, 019-19741)

### Image acquisition, processing, and quantitative analysis

Confocal microscopy was performed at the core facility bioimaging of the Biomedical Center (BMC) with an inverted Leica SP8 microscope using the LASX software (Leica). Overview images were acquired with a 10x/0.30 objective, higher magnification pictures with a 20x/0.75, 40x/1.30 or 63/1.40 objective, respectively. Images utilized for the microglia morphology analysis were acquired with an 40x/1.30 objective with an image matrix of 1024×1024 pixel, a pixel scaling of 0.2 μm x 0.2 μm and a depth of 8-bit. Image processing was performed using the NIH ImageJ software (version 2.1.0/1.53f). To acquire overview images, single images were stitched using the ImageJ plug-in tool pairwise stitching (Preibisch et al. 2009).

For all quantifications a minimum of two sections per animal were analyzed. In each section, an area of 300 μm (150μm on each side of the injury) was selected and either the pixel covered area or the number of positive cells in all individual z-planes of an optical z-stack was quantified. Additionally, to account for variations in section thickness, total cell numbers were normalized to the section depth. Statistical analysis was performed using GraphPad Prism (version 9.3.1).

### Spatial transcriptomics analysis

Mouse brains from 3dpi or intact *C57Bl6/J* mice were embedded and snap frozen in an isopentane and liquid nitrogen bath as recommended by 10x Genomics (Protocol: CG000240). During cryosectioning (Thermo Scientific CryoStar NX50) the brains were resected to generate a smaller sample (Fig. 1a) and two 10μm thick coronal sections of the dorsal brain area were collected in one capture area. The tissue was stained using H&E staining and imaged with the Carl Zeiss Axio Imager.M2m Microscope using 10x objective (Protocol: CG0001600). The libraries were prepared with Visium Spatial Gene Expression Reagent Kits (CG000239) with 18min permeabilization time and sequenced on an Illumina HiSeq1500 instrument and a paired-end flowcell (High output) according to manufacturer protocol, with sequencing depth of 55231 (Intact) and 75398 (3dpi) mean reads per spot. Sequencing was performed in the Laboratory for Functional Genome Analysis (LAFUGA).

Data were mapped against the mouse reference genome mm10 (GENCODE vM23/Ensembl 98; builds versions 1.2.0 and 2020A from 10xGenomics) with Space Ranger 1.2.2. Both datasets were analyzed, and quality checked following the Scanpy^68^ and Squidpy^69^ pipeline, selecting spots with at least 1500 reads and a maximum 45% mitochondrial fraction. Normalization and log transformation was performed using the counts per million (CPM) strategy with a target count depth of 10,000 using Scanpy’s^68^ normalize_total() and log1p functions. Following cell count normalization and scaling (function scale in Scanpy), experimental groups were integrated. Highly variable gene (HVGs) selection was performed via the function highly_variable_genes() using the Cell Ranger flavor with default parametrization, obtaining 2000 HVGs. Unsupervised clustering of cells was done using the Leiden algorithm^70^ as implemented in Scanpy. This allowed classification of multiple clusters based on marker genes selected using test_overestim_var() between the normalized counts of each marker gene in a cluster against all others (function rank_genes_groups in Scanpy). The layer marker score was performed using the function score_genes (as implemented in Scanpy) based on established marker genes (Ext. Table 3) described by Zeisel, A. et al 2018^33^. Gene ontology enrichment analysis was performed using the function enrichGO() (R package: clusterProfiler^52^) on the marker genes for cluster VI (indicated above) selecting the genes with pval<0.05 and log_2_fc>1 and the top 10 functions of the three aspects (MF: Molecular Function; CC: Cellular Component; BP: Biological Process) were presented on a dot plot.

### Single-cell analysis

The lesioned grey matter of the somatosensory cortex of C57BL/6J mice at 3dpi and 5dpi or the corresponding region of the noninjured cortex were isolated using a biopsy punch (∅ 0.25cm) and the cortical cells were dissociated at a single cell level using the Papain Dissociation System (Worthington, # LK003153) followed by the Dead Cell Removal kit (Miltenyi Biotec # 130-090-101), according to manufacturer’s instructions. Incubation with dissociating enzyme was performed for 60 min.

Single-cell suspensions were resuspended in 1xPBS with 0.04% BSA and processed using the Single-Cell 3’ Reagent Kits v2 or v3.1 (Ext. Table 13) from 10xGenomics according to the manufacturer instructions. In brief, this included generation of single cell gel beads in emulsion (GEMs), post-GEM-RT cleanup, cDNA amplification and library construction. Illumina sequencing libraries were sequenced on a HiSeq 4000 or NovaSeq6000 system (with an average read depth of 30,000 raw reads per cell) according to the manufacturer’s instructions for each version. Sequencing was performed in the genome analysis center of the Helmholtz Center Munich Transcriptome alignment of single-cell data was done using Cell Ranger v3.0.2 and v6.0.0 against the mouse reference genome mm10 (GENCODE vM23/Ensembl 98; builds versions 1.2.0 and 2020A from 10xGenomics). Quality Control (QC) of mapped cells from all datasets integrated was done using recommendations by Luecken and Theis^71^ selecting cells with at least 1000 genes, maximum of 50000 reads and 25% mitochondrial fraction. Doublets were removed using the Scrublet framework^72^, clusters expressing multiple lineage genes were identified as mixed population and were removed from the further analysis. Normalization was performed using the scran^68^ package (R package) followed by log-transformation using Scanpy’s log1p functions^68^. Highly variable gene (HVGs) selection was performed via the function highly_variable_genes using the Cell Ranger flavor with default parametrization, obtaining 2000 HVGs. Following cell count normalization and scaling, (function scale in Scanpy) experimental groups were batch corrected with scVI^73,74^. Unsupervised clustering of cells was done using the Leiden algorithm^70^ as implemented in Scanpy. This allowed classification of multiple main clusters based on marker genes selected using test_overestim_var between the normalized counts of each marker gene in a cluster against all others (function rank_genes_groups in Scanpy). The top 50 marker genes were used for the cluster annotation using the online available databases for the mouse brain (http://mousebrain.org) and the immune cells (http://rstats.immgen.org/MyGeneSet_New/index.html). Additionally, we generated gene expression scores using the function score_genes (as implemented in Scanpy) based on established marker genes (Table 2) of the main cell populations in the adult mouse brain to further confirm the cluster annotation. Visualization of cell groups is done using Uniform Manifold Approximation and Projection (UMAP)^75^, as implemented in Scanpy. Differential gene expression analysis between treated and control conditions was performed using the tool diffxpy (https://diffxpy.readthedocs.io/en/latest/index.html) using the Wald test. Of note, since some glial subclusters are comprised of only few cells, the differential gene analysis did not reveal differential expressed genes in these subclusters.

All the comparisons of the overlapping genes were performed using the R package UpSetR^76^ which provides an efficient way to visualize the intersecting gene set in UpSet plot. For cluster comparison Additionally, the gene ontology analysis was performed using the R package clusterProfile^52^, using the functions compareCluster (fun:enrichGO) or enrichGO. The visualisation of the functional enrichment results was done using the following visualization methods from the R package enrichplot^52^: dot plot; enrichment map (function: emaplot) (based on the pairwise similarities of the enriched terms calculated by the pairwise_termsim function); and the Gene-Concept Network plot (function: cnetplot).

### Spatial alignment of the scRNA-seq data

For the spatial localization of the scRNA-seq data, we used the Python package Tangram^46^, focusing on the 3dpi control condition and using only the cortical cluster of the Visium dataset in order to have the same anatomical region. We selected the training genes using the tool AutoGeneS^77^ and used 439 training genes as the union of the top informative marker genes of each cluster in the scRNA-seq data that were detected in the Visium profiles. To find the spatial alignment for the scRNA-seq we used the Tangram^46^ function map_cells_to_space() which gave us the probabilistic mapping score. Additionally, we segmented the H&E image, using the Squidpy^69^ function segment which was used for deconvolving the Visium data using the Tangram^46^ functions count_cell_annotations() and deconvolve_cell_annotations().

### Spatial gradient analysis

Spatial gradients extending from the lesion core towards perilesional regions were defined using SPATA^78^ and its successor SPATA2^39^ (under development; https://themilolab.github.io/SPATA2/). The Scanpy-processed object described above was used as input. Both lesion cores were manually annotated based on the H&E staining using createImageAnnotations(). Visium spots were binned into concentric circles using the following arguments: n_bins_circle = 13, binwidth = “95μm”. Spots from non-cortical clusters (III,V,X,XIII,XIV,XV,XII,XVI) were excluded from the analysis using the argument bcsp_exclude. Genes with >50 total counts were screened for their correlation with pre-defined gradients (*e*.*g*. linear descending) using imageAnnotationScreening(), for both injuries separately. Spot metadata derived from Scanpy and Tangram, as well as genes that correlated most strongly with selected pre-defined gradients (sorted by p_value_mean) were plotted using plotIasHeatmap_merge() and plotIasRidgeplot_merge(), custom adaptations of original SPATA2 functions, in which values represent the bin-wise mean of both injuries. Descending models included ‘linear_descending’, ‘immediate_descending’, ‘abrupt_descending’, ‘late_descending’. Ascending models included ‘linear_ascending’, ‘immediate_ascending’, ‘abrupt_ascending’, ‘late_ascending’ (see function showModels()). For screening of gene sets, the following sets were downloaded from MsigDB (https://www.gsea-msigdb.org/gsea/msigdb/mouse/collections.jsp): Biocarta, KEGG, Reactome, WikiPathways, GO (MF/CC/BP), Hallmark. Per gene set, the mean expression of all included genes was calculated and screened for correlation with the same pre-defined gradients as described for single genes. A snapshot of the utilized state of SPATA2 including custom functions is available at https://github.com/simonmfr/SPATA2/tree/publicationCK.

## Data availability

All sequencing data generated in association with this study are available in the Gene Expression Omnibus as a SuperSeries. Access can be provided upon request.

Details of analysis pipeline libraries are listed in Methods and available at https://github.com/NinkovicLab/Koupourtidou-Schwarz-et-al (private repository). A public repository will be created as soon as the manuscript is published. Notebooks and all files in the repository can be provided upon request.

