## Extended Figures for "Shared inflammatory glial cell signature after brain injury, revealed by spatial, temporal and cell-type-specific profiling of the murine cerebral cortex"

Koupourtidou, Schwarz, et. al. ED_Fig. 1

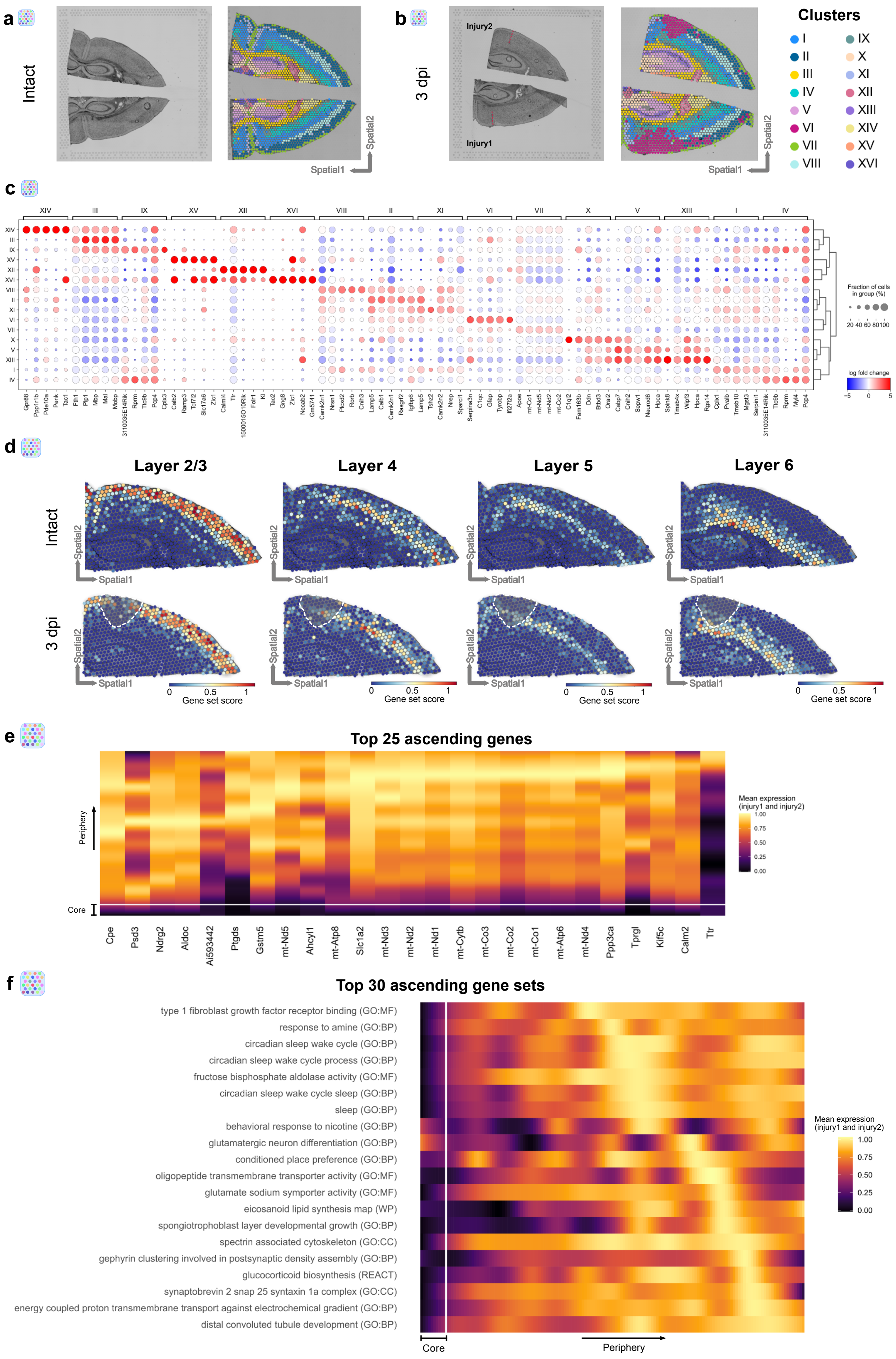

Koupourtidou, Schwarz, et. al. ED_Fig. 2

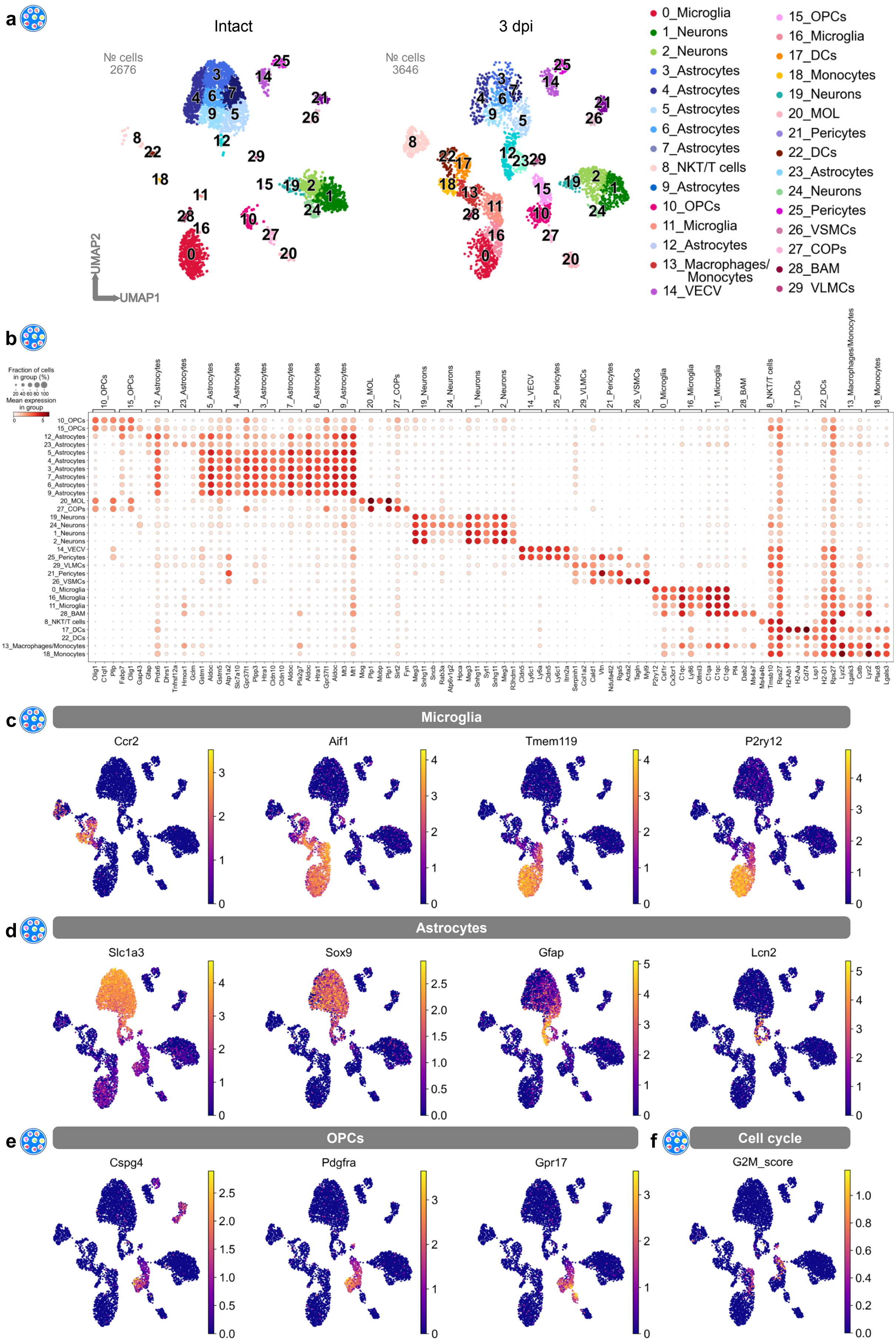

Koupourtidou, Schwarz, et. al. ED_Fig. 3

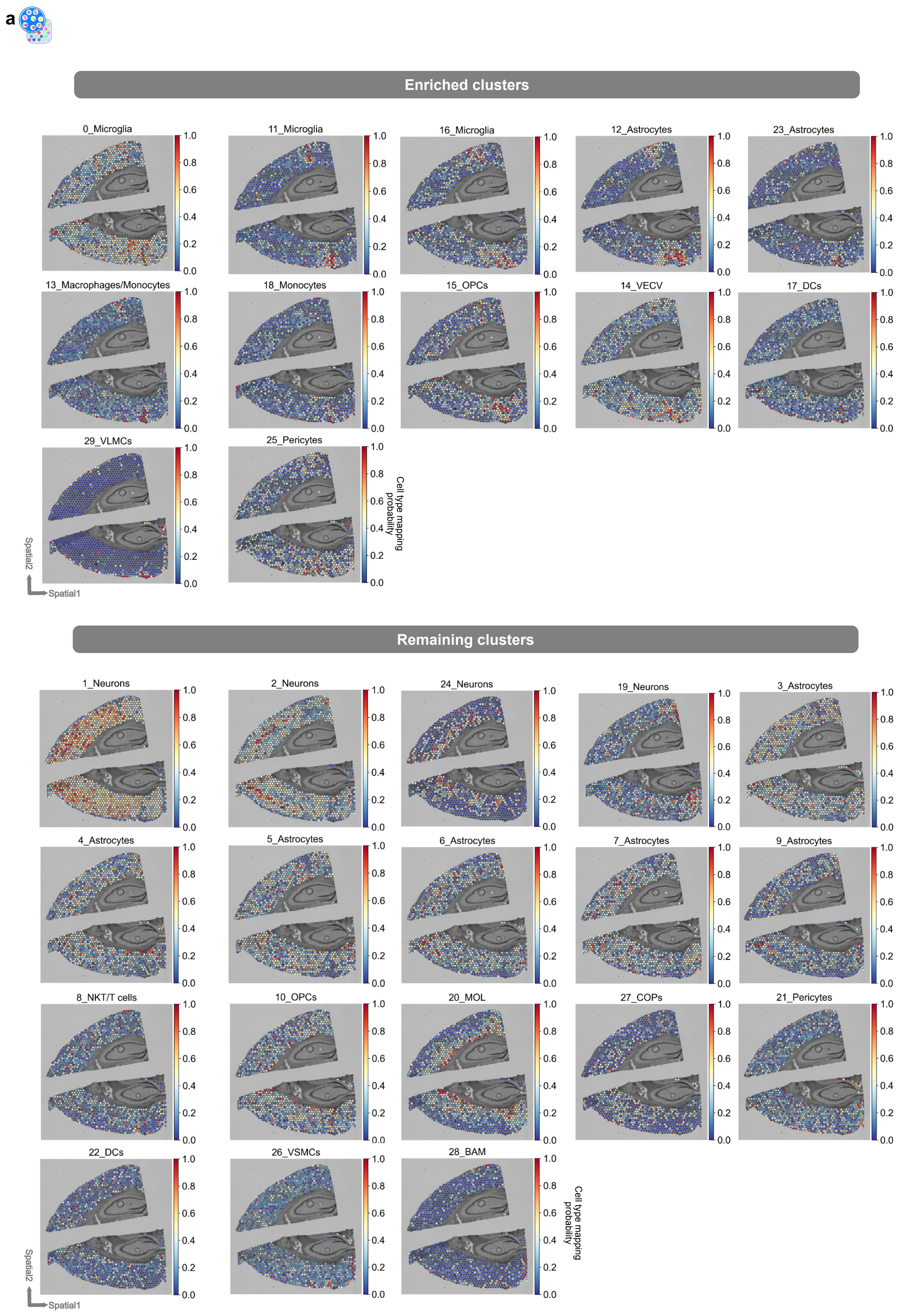

Koupourtidou, Schwarz, et. al. ED_Fig. 4

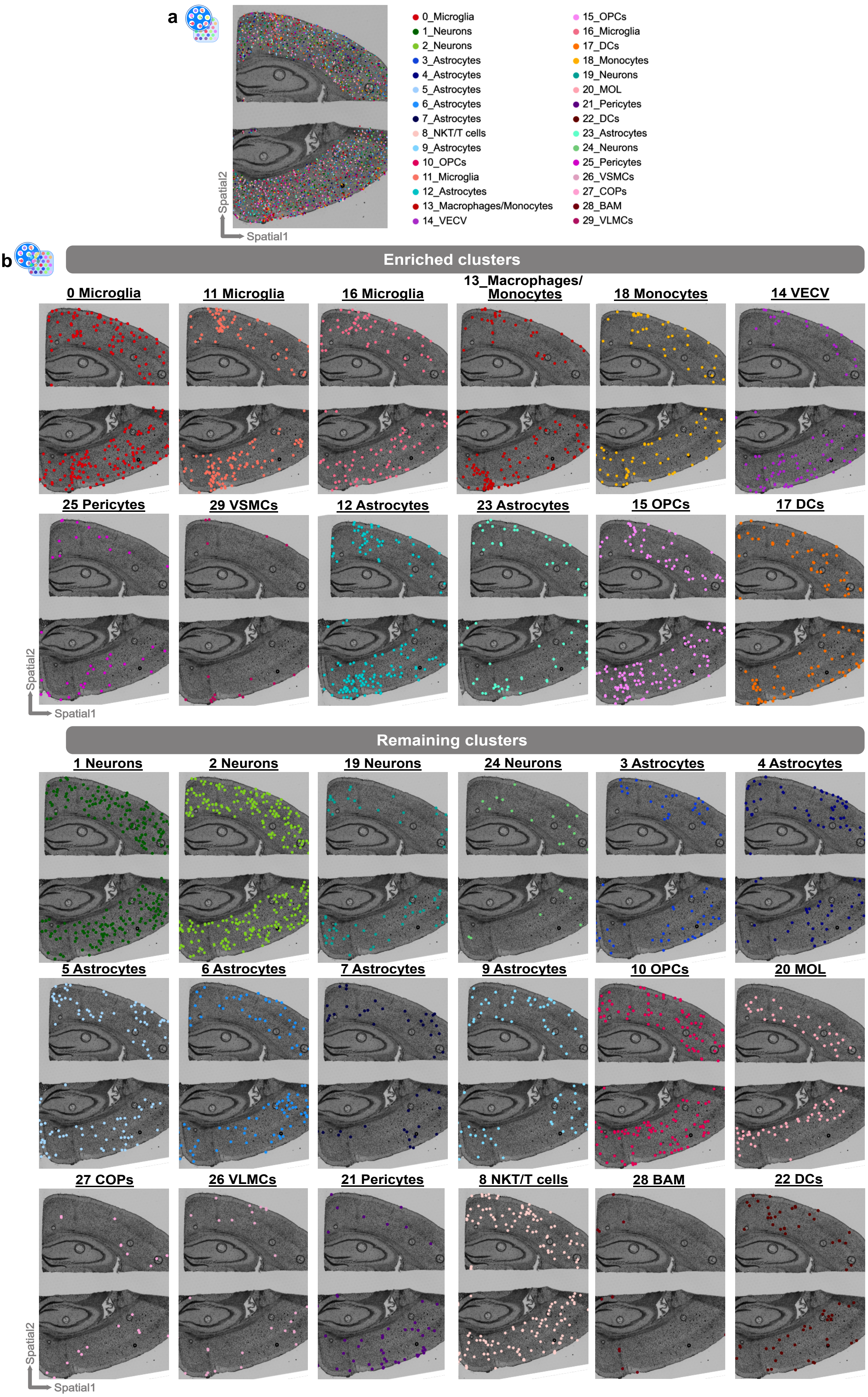

Koupourtidou, Schwarz, et. al. ED_Fig. 5

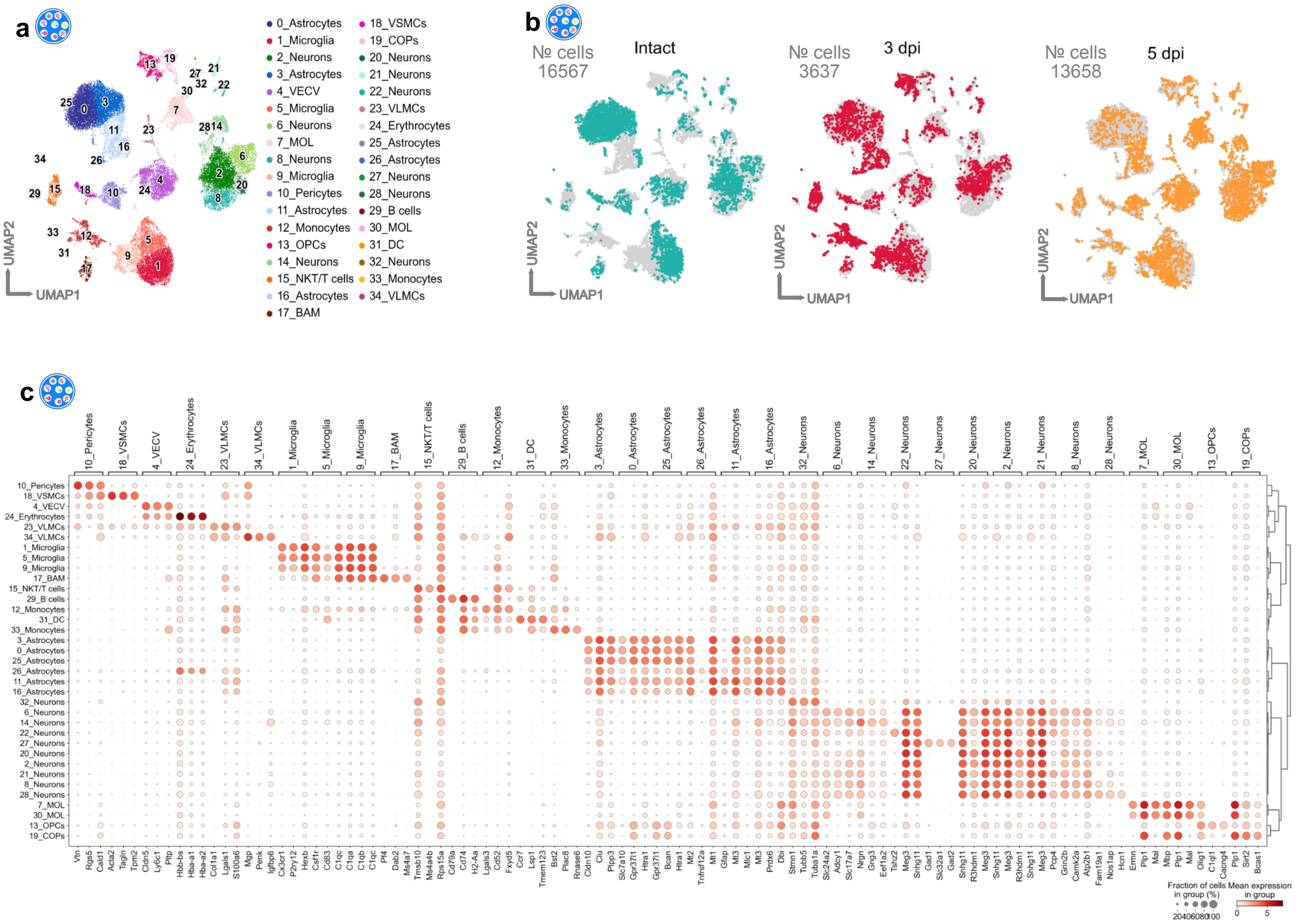

Koupourtidou, Schwarz, et. al. ED_Fig. 6

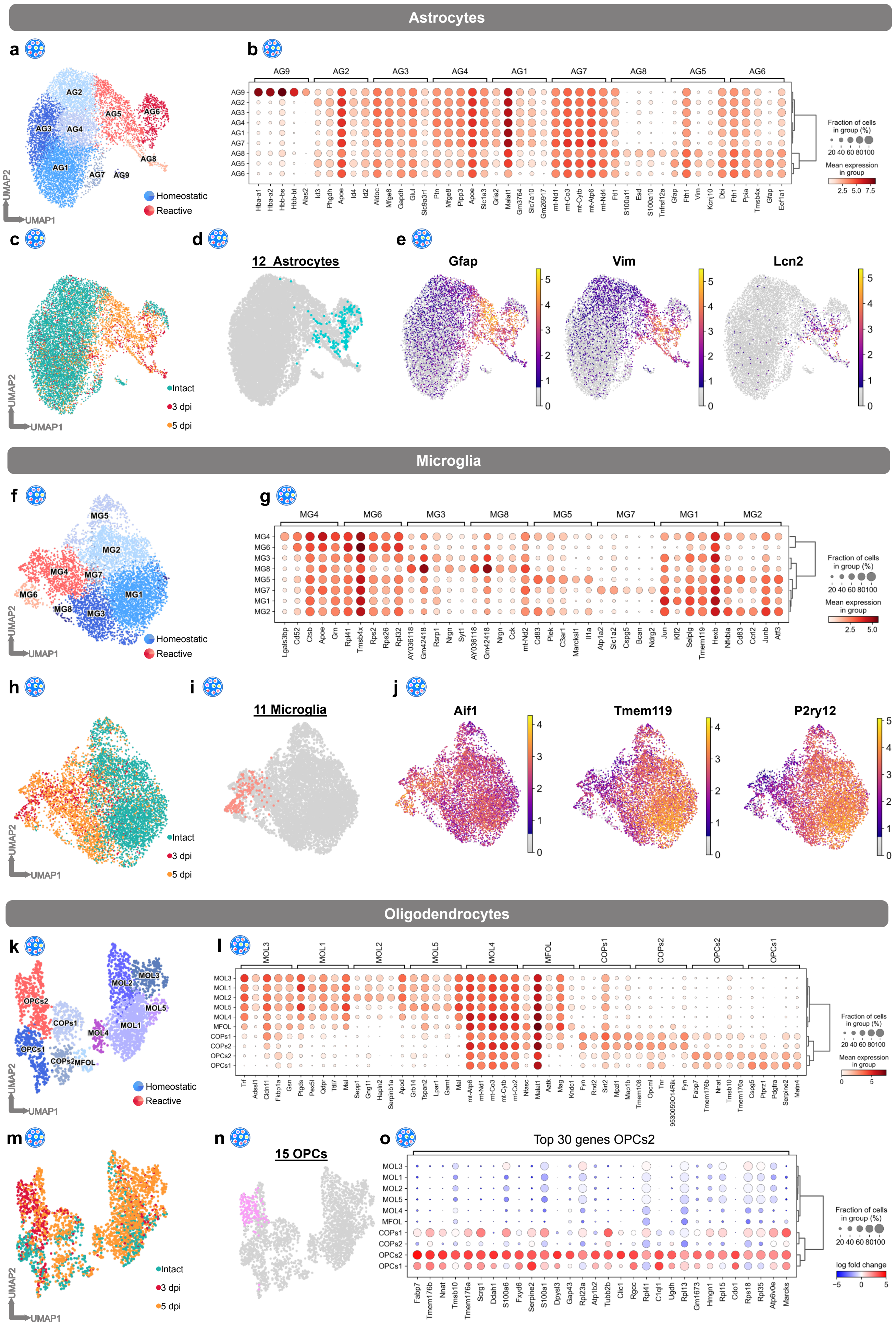

Koupourtidou, Schwarz, et. al. ED_Fig. 7

a

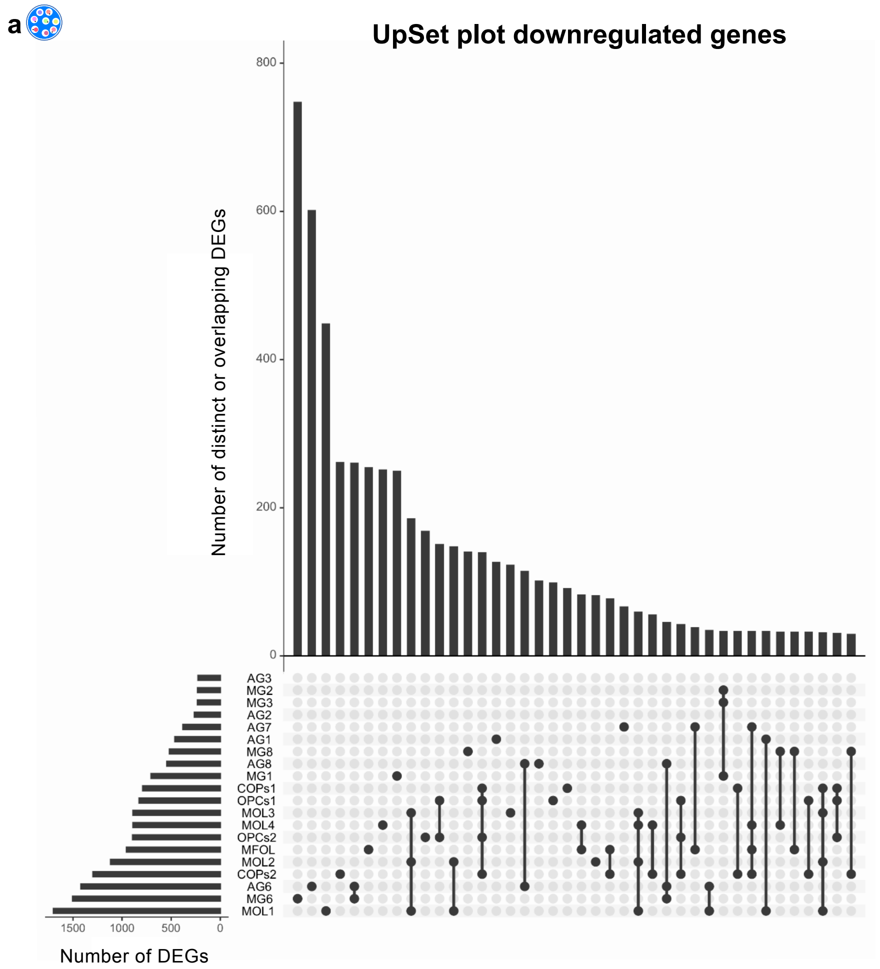


b

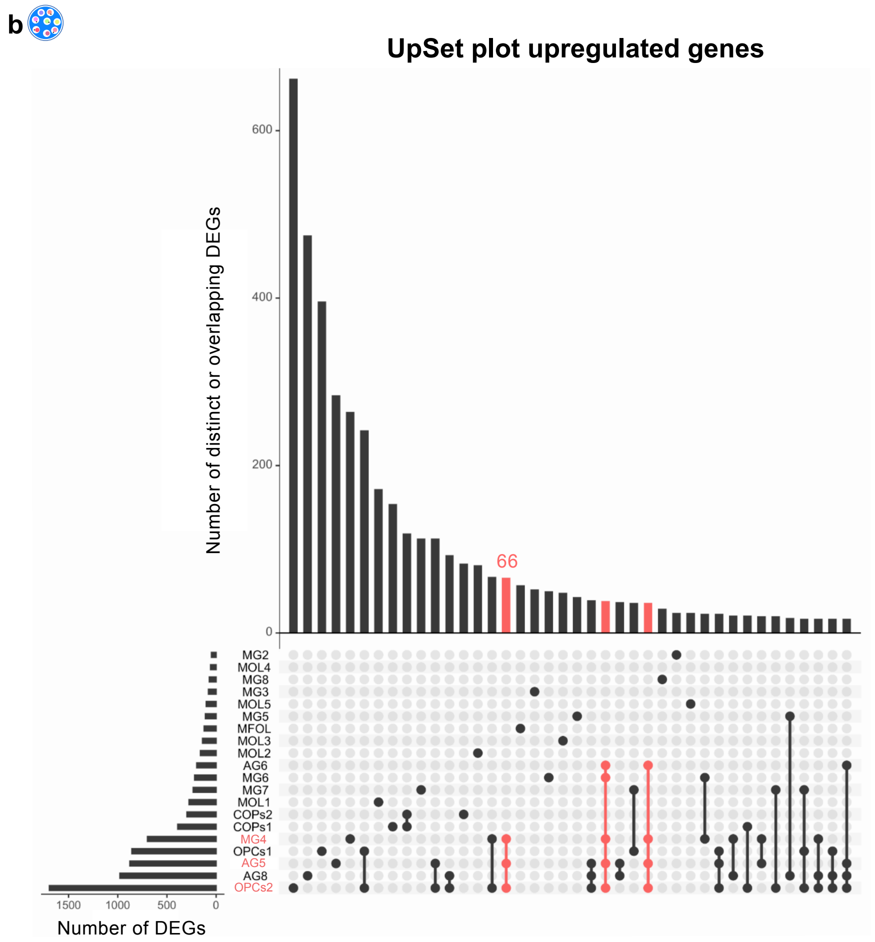

Koupourtidou, Schwarz, et. al. ED_Fig. 8

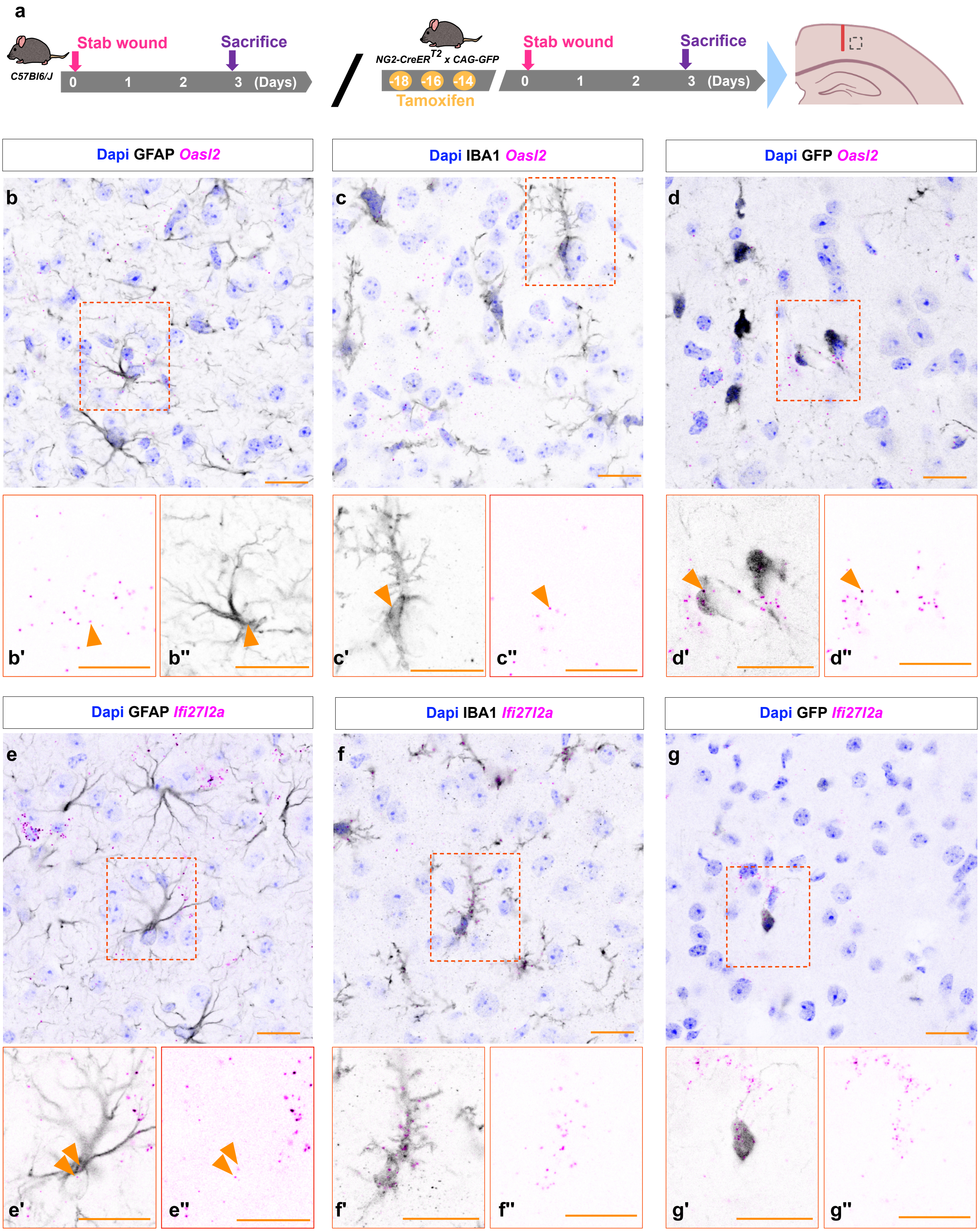

Koupourtidou, Schwarz, et. al. ED_Fig. 9

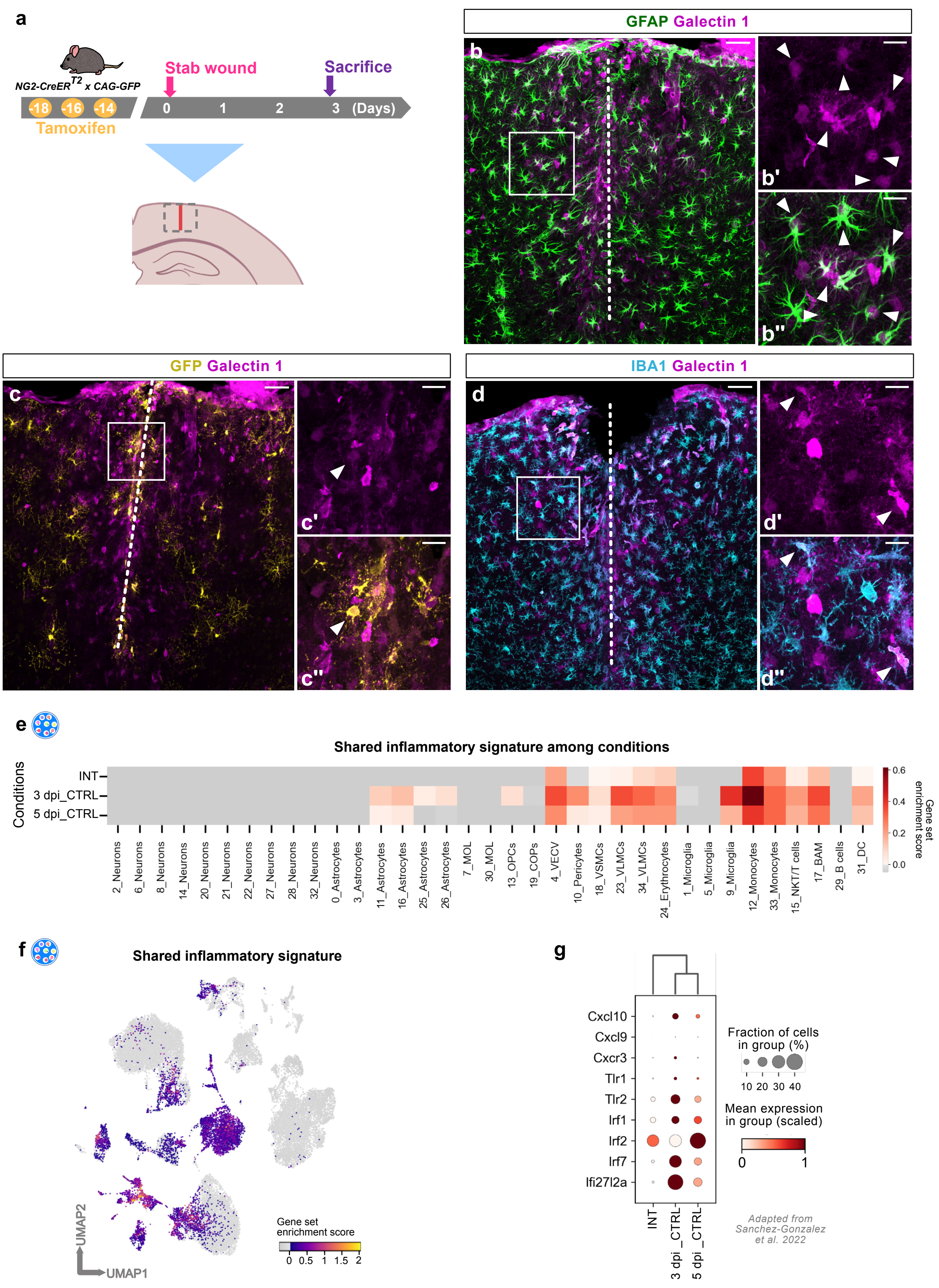

Koupourtidou, Schwarz, et. al. ED_Fig.10

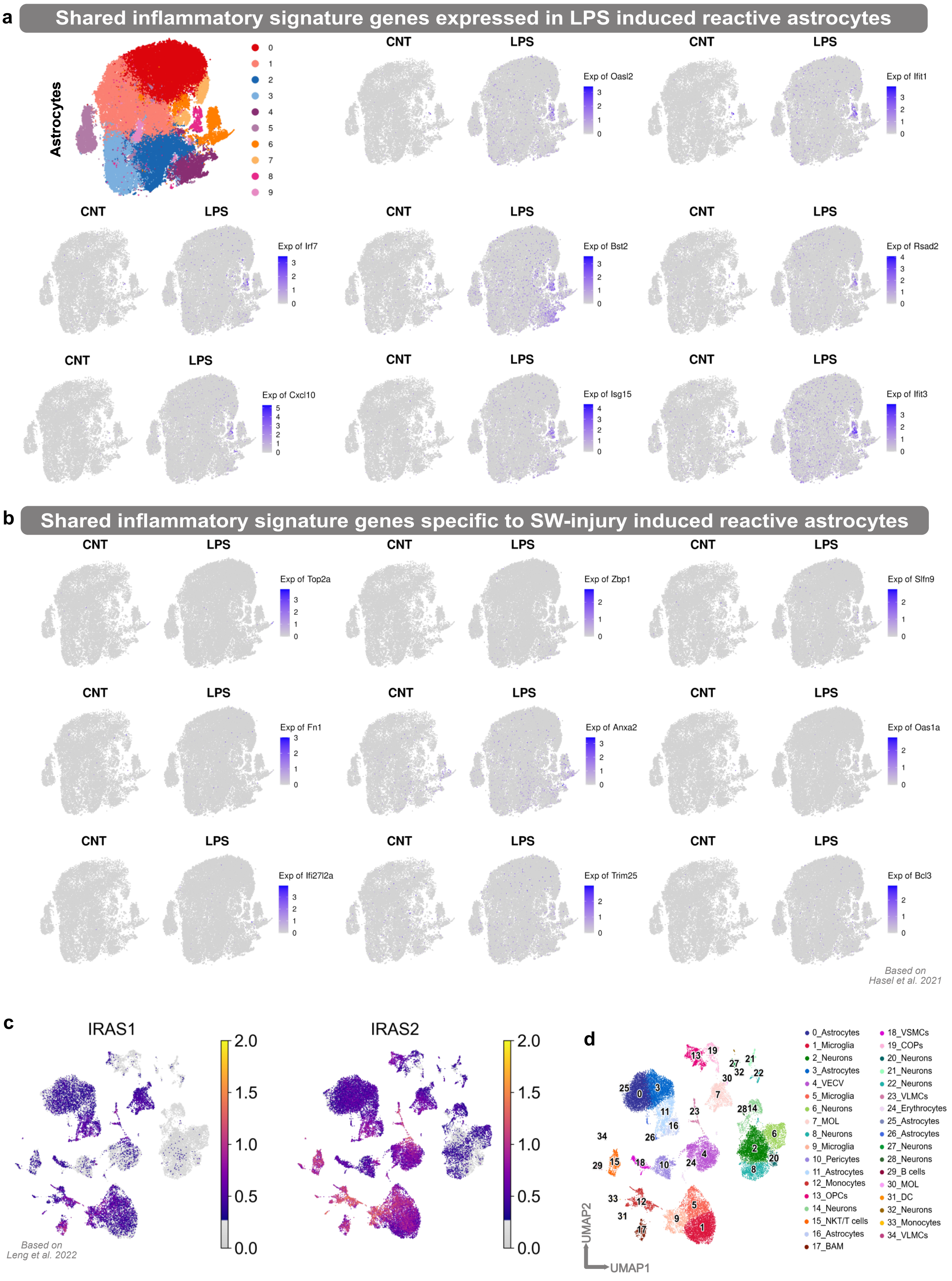

Koupourtidou, Schwarz, et. al. ED_Fig. 11

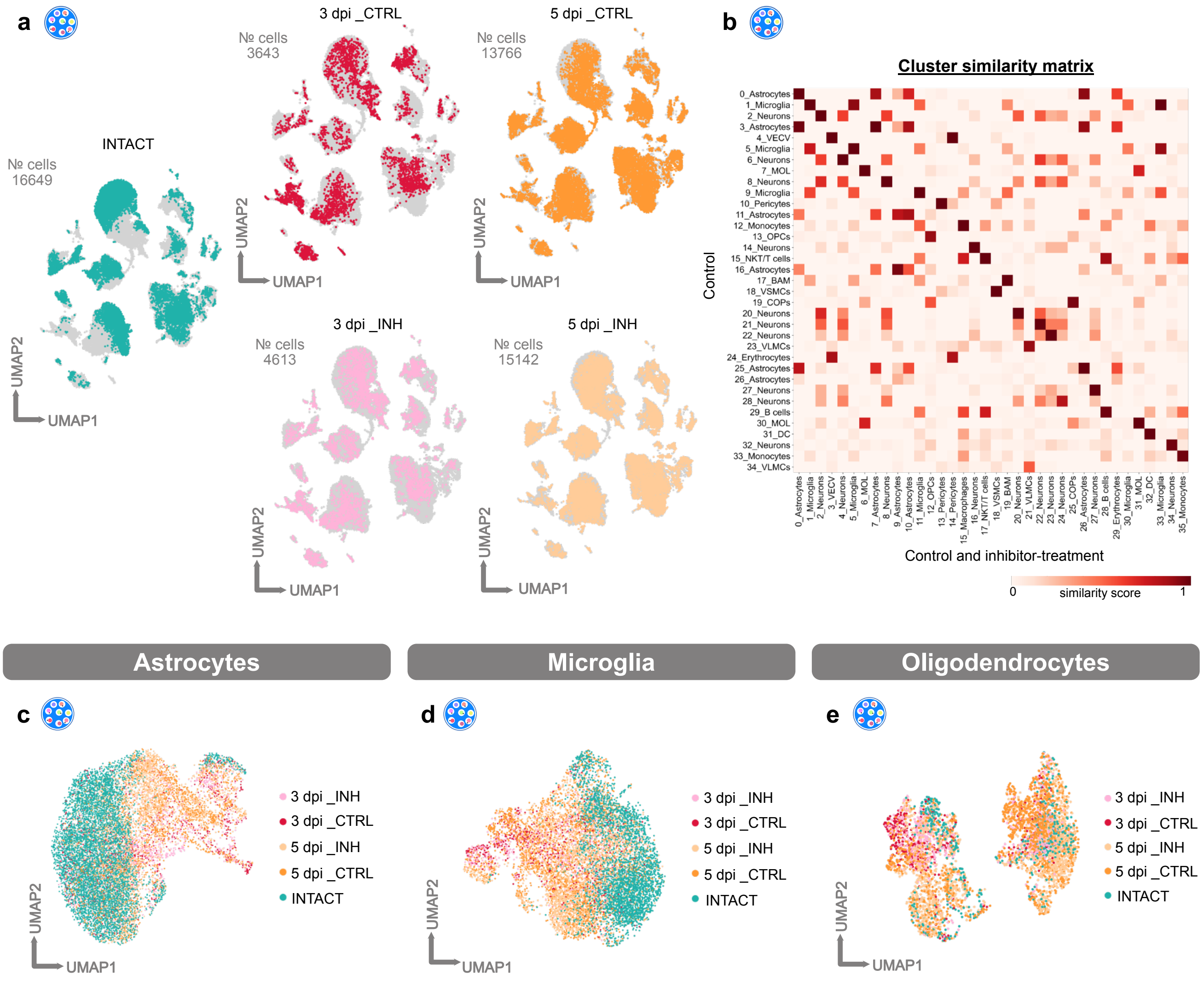

Koupourtidou, Schwarz, et. al. ED_Fig. 12

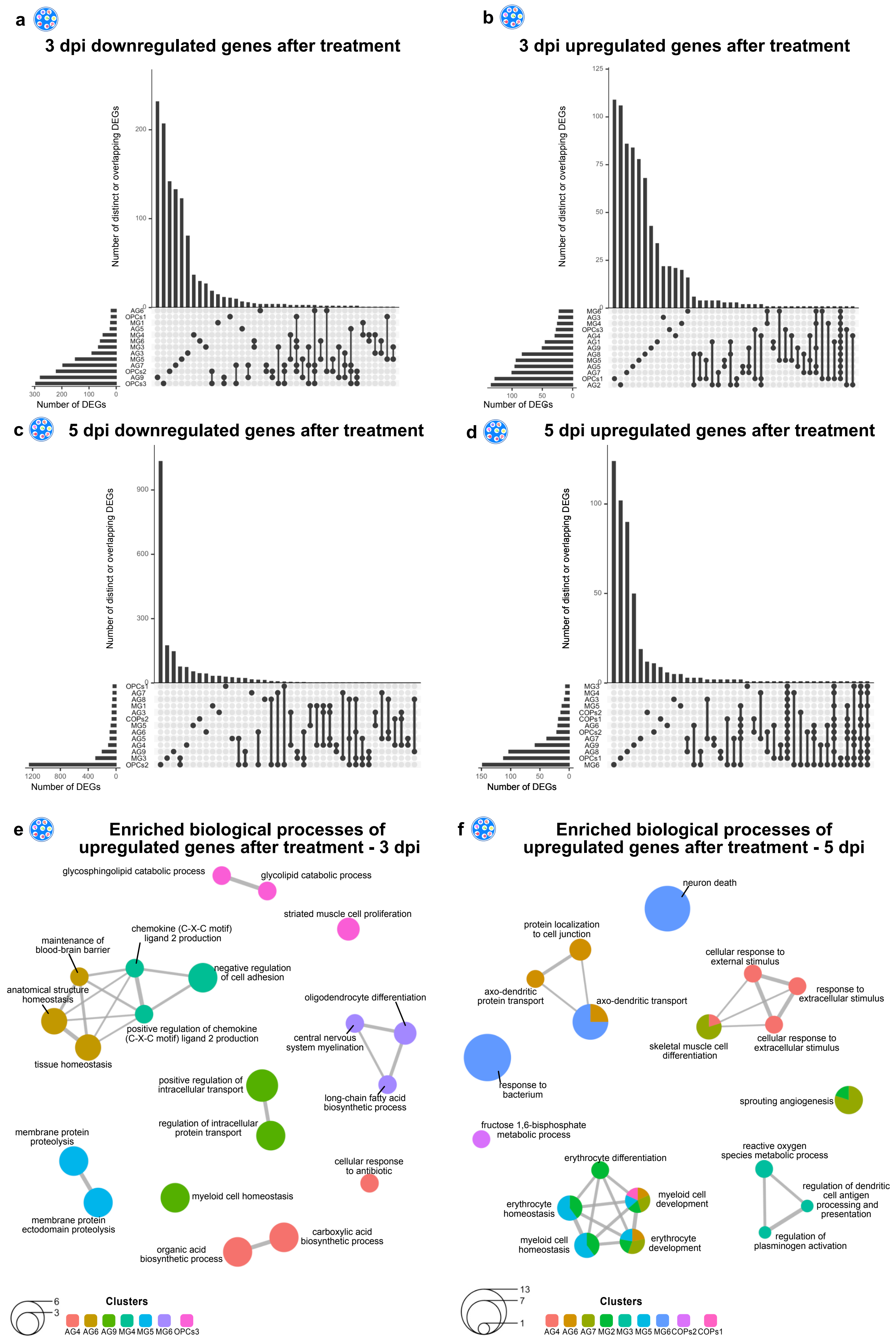

Koupourtidou, Schwarz, et. al. ED_Fig. 13

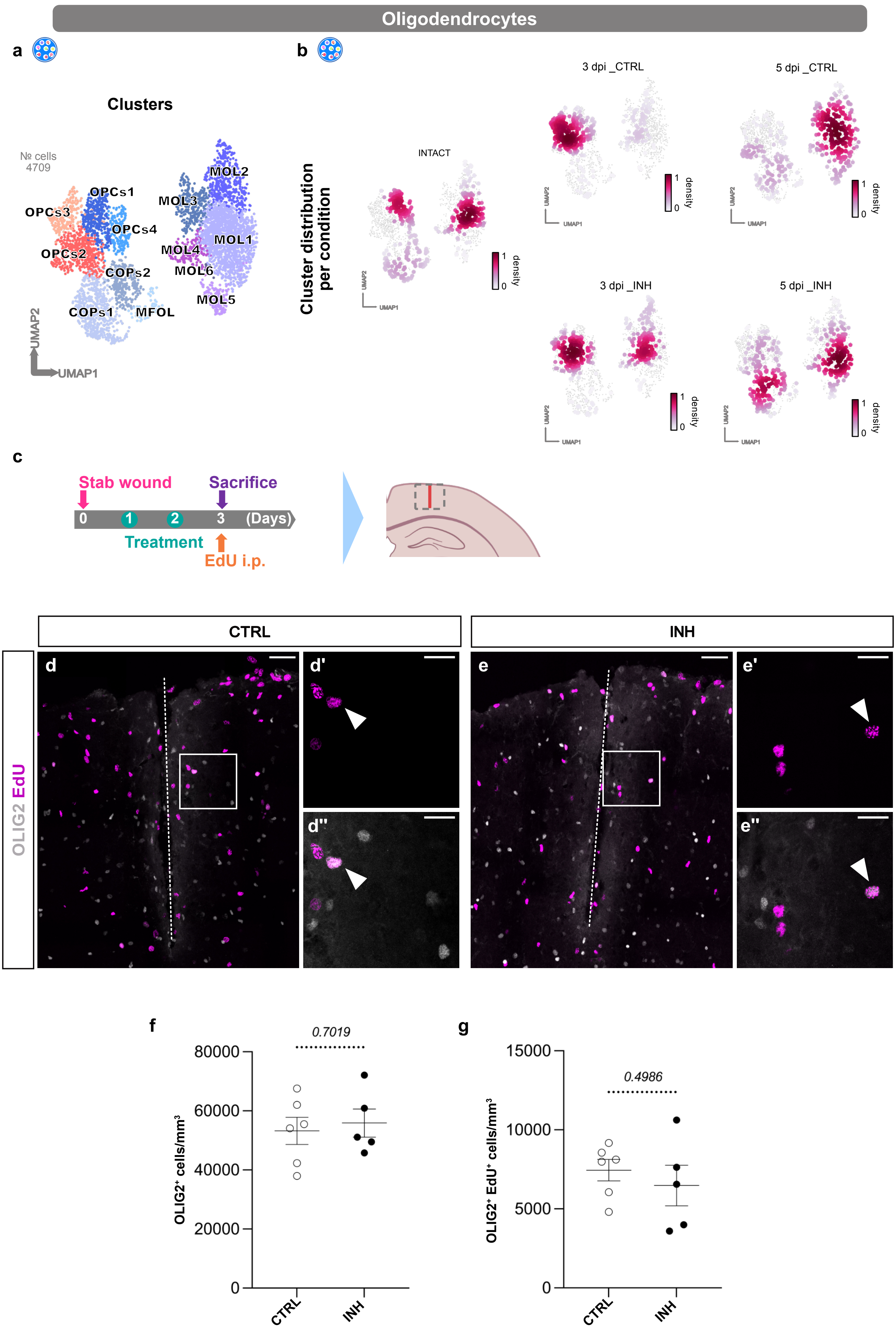

Koupourtidou, Schwarz, et. al. ED_Fig. 14

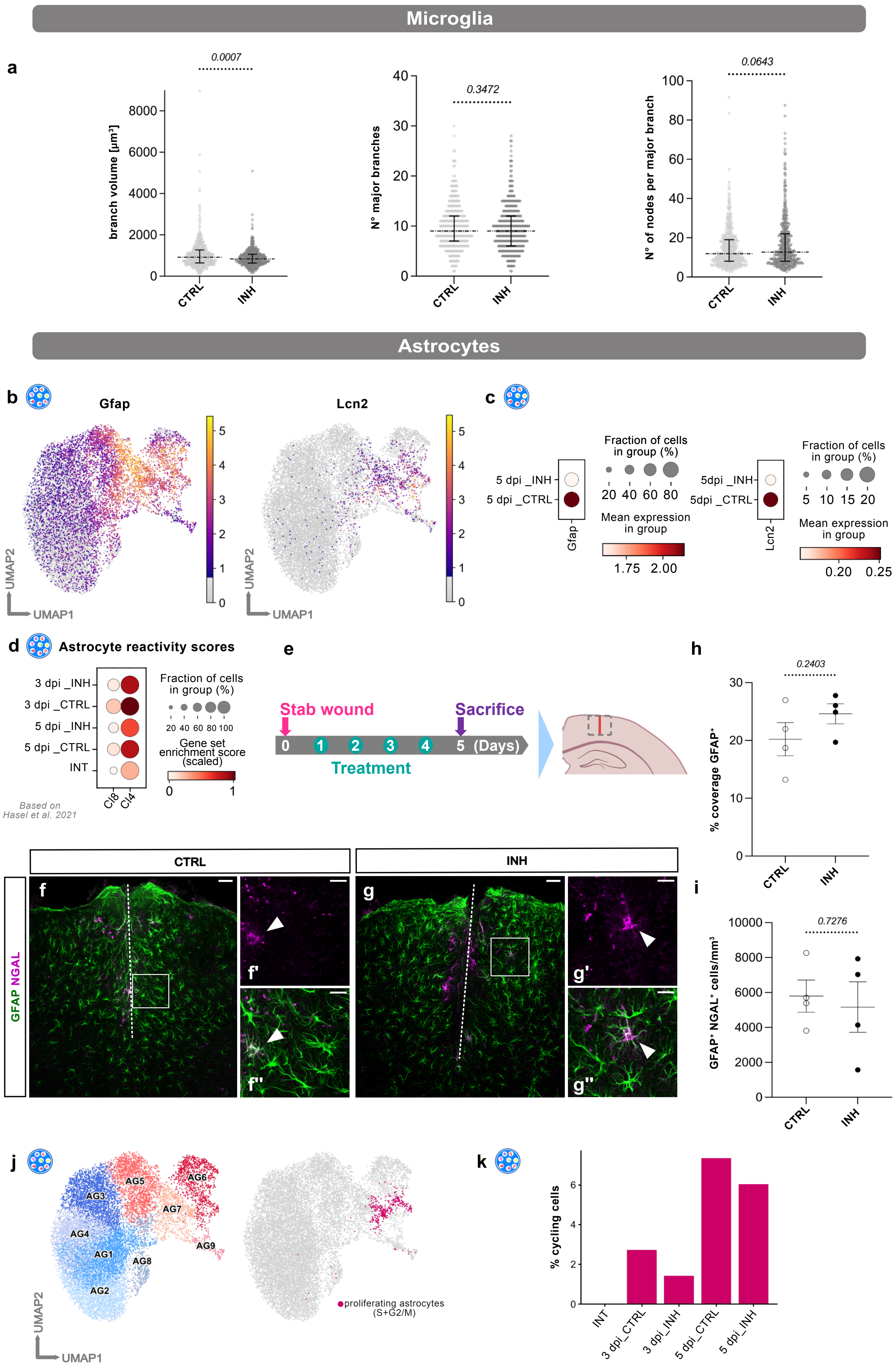
